# Chemogenetic Pericyte Activation Reveals Broad Contractile Ability and Limbic Vulnerability to Capillary Flow Deficits

**DOI:** 10.64898/2026.04.11.717917

**Authors:** Liam T. Sullivan, Dickson T. Chen, Catherine Foster, Ben Zimmerman, Kevin Elk, Bradley Marxmiller, Taylor McGillis, Yuandong Li, Stephanie K. Bonney, Lila Faulhaber, Dimitrios Davalos, Juliane Gust, Zhen Zhao, Anusha Mishra, Andy Y. Shih

## Abstract

Capillary pericytes contact most of the brain’s microvasculature. Yet, their influence on blood flow remains incompletely understood both locally in capillary networks and across brain regions. Here, we sparsely expressed the chemogenetic actuator Gq-DREADD in mouse brain pericytes to stochastically probe their contractility and influence on tissue perfusion and oxygenation. Chemogenetic stimulation induced robust contraction of pericytes over minutes, including those with thin processes on capillary segments proximal to venules, indicating control of vascular tone across the entire capillary bed. Pericyte contraction combined circumferential compression with longitudinal tension applied through thin processes, resulting in vasoconstriction and hypoperfusion. Histology revealed hypoxic microdomains induced by sparse pericyte contraction across the brain, and enriched hypoxia in limbic structures including the amygdala and hippocampus. These microscale perfusion deficits were undetectable by arterial spin labeling MRI. Our findings reveal the *in vivo* mechanics of pericyte contraction and identify limbic system vulnerability to capillary flow impairment.

## INTRODUCTION

The brain is one of the most vascularized organs in the body, with complex capillary networks that efficiently and dynamically supply oxygen and nutrients to support neural activity.^1,2^ The majority of this vasculature consists of dense capillary networks, where blood cells course, single-file, through the vessel lumen. The diameter of the capillary lumen is fine-tuned for proper distribution of blood within the network, and deviations from the normal diameter can result in blood flow maldistribution, capillary regression, and impaired metabolic supply.^3–5^ Impaired capillary perfusion is increasingly recognized as a basis for cerebral hypoperfusion in a range of neurological diseases, including Alzheimer’s disease^6^, stroke^7^ and epilepsy^8^. However, there remains limited knowledge on how vulnerabilities in capillary perfusion arise during disease, both locally within capillary networks and across distinct brain regions.

Pericytes are mural cells that line the walls of brain capillaries and have been strongly implicated in cerebral blood flow (CBF) impairment during disease.^9–14^ Pericytes possess a range of morphologies dependent on their location in the capillary network.^13^ Those located closer to the penetrating arteriole, termed ensheathing pericytes of the arteriole-capillary transition (ACT) zone, have circumferential processes, express the key contractile protein α-smooth muscle actin (α-SMA), and control dynamic second-to-second regulation of blood flow into the capillary bed. In contrast, pericytes further downstream express undetectable levels of α-SMA and possess long, thin processes that trail like vines along the endothelium. These “thin-strand pericytes” represent the majority of pericytes and their spindly morphology and deficiency in α-SMA has raised skepticism on their ability to regulate vessel tone.^9,15^ Prior studies shed some light on this debate by optogenetically stimulating thin-strand pericytes to show that they could indeed contract, albeit with much slower kinetics than arteriolar smooth muscle cells.^3,16–19^ However, many questions remained, including how the morphology of thin-strand pericytes could confer contractile force, and if all pericytes from arteriole to venular poles of the capillary bed possessed contractile capabilities. Further, it was unclear whether pericytes beyond the cerebral cortex were contractile and able to influence tissue oxygen, given marked variations in capillary density and architecture across brain regions.^1,2,20^ Technical limitations hampered progress. Among these limitations, optogenetic contraction of pericytes required prolonged stimulations that were challenging to combine with imaging, sensitive to phototoxicity, and limited to the superficial cortex. Further, prior genetic targeting tools for pericytes were neither brain-nor pericyte-specific. This impasse left fundamental questions on pericyte roles in cerebral blood flow control.

Recently, the advent of an inducible central nervous system (CNS) pericyte-targeting Cre mouse line (Atp13a5-CreERT^2^-IRES-tdTomato)^21^ paved the way for a new strategy - pericytes chemogenetics.^22,23^ Pericytes express high levels of Gq-dependent G-protein coupled receptors (Gq-GPCRs), such as thromboxane A2 receptor and endothelin-1 Type A receptor, that promote vasoconstrictive signaling.^24^ The action of these receptors can be mimicked through pericyte-specific expression of the chemogenetic actuator, Gq-DREADD.^11^ We leveraged this approach to sparsely express Gq-DREADD in capillary pericytes, enabling their controlled contraction *in vivo* following systemic administration of a blood-brain barrier (BBB) permeable DREADD-specific ligand. Sparse expression allowed pericyte contractile responses to be cleanly isolated from the influence from upstream and neighboring mural cells. The stochastic nature of genetic targeting also allowed pericytes to be examined in different regions of the capillary network. Finally, histology could be used to examine brain-wide effects on tissue oxygenation. Together, these advances reveal that pericytes exert potent control over capillary perfusion across the brain.

## RESULTS

### Genetic targeting of pericytes in the CNS

To chemogenetically target CNS pericytes, we used the recently developed Atp13a5-CreERT2-IRES-tdTomato mouse line, herein referred to as Atp13a5-CreERT2 mice.^21^ This line expresses both CreERT2 and tdTomato for conditional genetic manipulation and optical detection of brain pericytes, respectively. To confirm cell specificity, we performed *in vivo* two-photon imaging in the somatosensory cortex and confirmed tdTomato expression in pericytes across the capillary bed, but lack of labeling in arteriolar smooth muscle cells (SMCs) of pial or cortical penetrating arterioles (**Supplementary Fig. 1A-B**). Expression was also observed in mural cells of ascending venules, but minimal expression on pial venules. Additionally, histological analyses indicated tdTomato expression in >95% of PDGFRβ+ cells along capillary-sized vessels, further supporting CNS pericyte targeting specificity in the brain, spinal cord and retina (**Supplementary Fig. 1C,D, Supplementary Fig. 2)**.

To evaluate the efficiency and specificity of Cre-induced recombination, we crossed Atp13a5-CreERT2 mice with conditional reporter mice (Ai3-YFPfl/fl), which express YFP upon tamoxifen induction (**Fig 1A, Supplementary Fig. 1E,F, Supplementary Fig. 3**). Histological analyses indicated ∼50-70% recombination efficiency across brain regions, with slightly lower levels in the spinal cord (**Fig 1B**). Recombination in retinal pericytes was lowest at ∼10%. Critically, both tdTomato and Cre-induced YFP reporter expression were absent in a broad range of peripheral tissues examined, again consistent with CNS pericyte specificity (**Supplementary Fig. 1F, Supplementary Fig. 4**).

**Figure 1.**
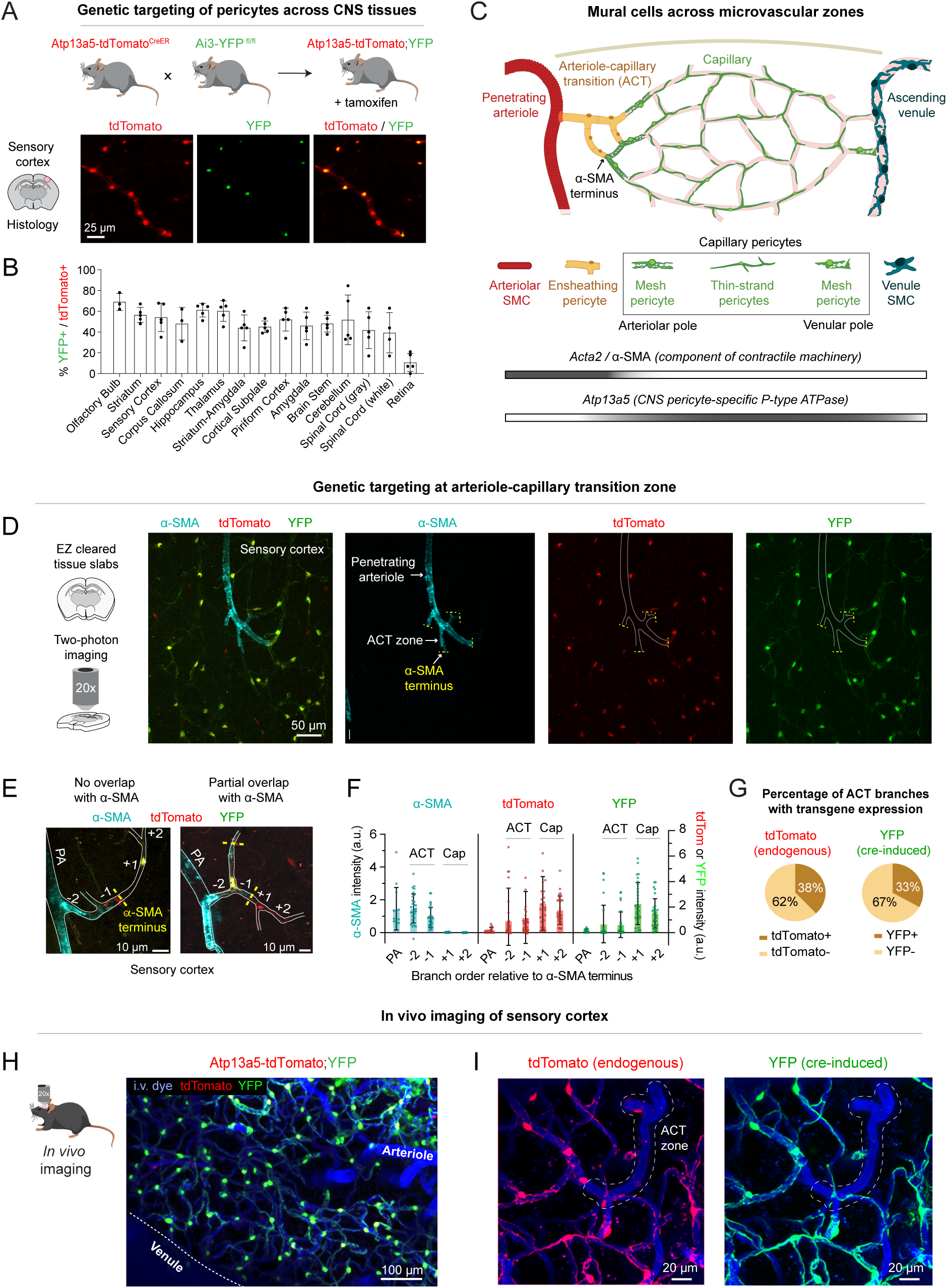
Genetic targeting of CNS pericytes using Atp13a5-CreERT2 mice. **(A)** Atp13a5-CreERT2 mice were crossbred with Ai3 (YFP) reporter mice to evaluate specificity and penetrance of Cre-dependent gene expression in CNS pericytes using histology. **(B)** Percentage of pericytes with YFP reporter expression across CNS regions. **(C)** Schematic showing mural cell subtypes across microvascular zones. *Acta2* and *Atp13a5* gene expression is compared along the arteriovenous axis. **(D)** Optically-cleared brain tissue slabs from Atp13a5-CreERT2;Ai3 mice immunostained for α-SMA and imaged by two-photon to evaluate fluorescent protein expression in relation to the ACT zone. **(E)** Fluorescent reporter expression as a function of branch order from α-SMA terminus (dotted yellow line). Note that the pericyte just before α-SMA terminus occasionally has lower but discernable α-SMA expression compared to mural cell further upstream. **(F)** Bar graph showing reporter intensity at the penetrating arteriole (PA) and different branch orders from the α-SMA terminus. Negative branch orders correspond to the ACT zone, covered by α-SMA-positive ensheathing pericytes. Positive branch orders correspond to the capillary zone, covered by α-SMA-low/undetectable capillary pericytes. **(G)** Percentage of ACT zone branches that express Atp13a5 and are genetically targeted. Total 148 ACT vessels examined in cortex (58 vessels), thalamus (51 vessels) and hippocampus (39 vessels) across 3 mice. **(H)** *In vivo* two-photon imaging of Atp13a5-CreERT2;Ai3 mouse sensory cortex through a chronic cranial window, showing robust YFP labeling in pericytes but not mural cells of pial/penetrating arterioles or pial venules. **(I)** High-magnification image of sensory cortex showing estimated ACT zone and surrounding capillary networks.

Brain vascular single-cell transcriptomics indicate that *Atp13a5* is expressed mostly by capillary pericytes, which have undetectable levels of α-SMA in the brain (**Fig. 1C**).^25^ However, it was unclear if ensheathing pericytes in the ACT zone upstream were also genetically targeted. The shift from ensheathing pericytes to downstream capillary pericytes is typically marked by a sharp termination in α-SMA expression and transition from circumferential to mesh or thin longitudinal processes.^13^ Capillary pericyte are comprised primarily of thin-strand pericytes, but pericytes with mesh-like processes can reside at the arteriole and venular poles of the capillary bed. To determine the extent of ensheathing pericyte targeting, we imaged thick coronal brain sections from Atp13a5-CreERT2:Ai3-YFP^fl/fl^ mice that were immunostained for α-SMA and optically cleared *for post-mortem* two-photon imaging (**Fig. 1D**). In imaged tissue volumes, we located α-SMA termini on branches of penetrating arterioles in the somatosensory cortex, thalamus and hippocampus. Microvessel segments were then counted upstream and downstream from this point to measure the extent of YFP expression in ensheathing pericytes (upstream) vs. capillary pericytes (downstream), respectively (**Fig. 1E-F**). This revealed that a sub-population of ensheathing pericytes were genetically targeted (**Fig. 1G, Supplementary Fig. 5**). To better differentiate ensheathing pericytes, we analyzed the intensity of co-expressed α-SMA and tdTomato/YFP and determined that 33% of ensheathing pericytes were genetically targeted in brain regions examined, including cortex, hippocampus and thalamus (**Supplementary Fig. 6**). Thus, Atp13a5-CreERT2 mice primarily target capillary pericytes and some ensheathing pericytes gating flow into the capillary bed, but not SMCs of penetrating or pial arterioles.

*In vivo* two-photon imaging of the somatosensory cortex also identified robust YFP labeling of pericytes throughout the microvasculature of the somatosensory cortex, but not SMCs along pial and penetrating arterioles, again consistent with preferential targeting of capillary pericytes (**Fig. 1H,I)**. Critically, no parenchymal cell types were seen *in vivo*, However, some venous SMCs and fibroblast-like cells within the meninges were found to express tdTomato in Atp13a5-CreERT2 mice (**Supplementary Fig. 7**).

### Chemogenetic activation of CNS pericytes results in potent vasoconstriction

Having characterized CNS pericyte targeting specificity, we next established mice for chemogenetic stimulation *in vivo*. Atp13a5-CreERT2 mice were bred with mice carrying one allele of floxed Gq-DREADD (hM3Dq). The bigenic progeny will herein be referred to as Atp13a5-Gq-DREADD (**Fig. 2A**). Unlike the floxed Ai3 YFP reporter, the Gq-DREADD transgene was not inserted into the ROSA26 locus as intended during the production of these mice (verified by Jackson Labs), and in practice, we observed much lower efficiency of Cre recombination. Consequently, this provided sparse labeling of pericytes with the same tamoxifen regimen used for the Ai3 reporter. On average, only 4.6% (1-9% range) of total pericytes expressed Gq-DREADD in the somatosensory cortex, based on immunostaining for the HA-tag in the Gq-DREADD protein (**Supplementary Fig. 8A,B**). We leveraged this sparsity in expression for stochastic assessment of individual pericyte responses across the capillary bed.

**Figure 2.**
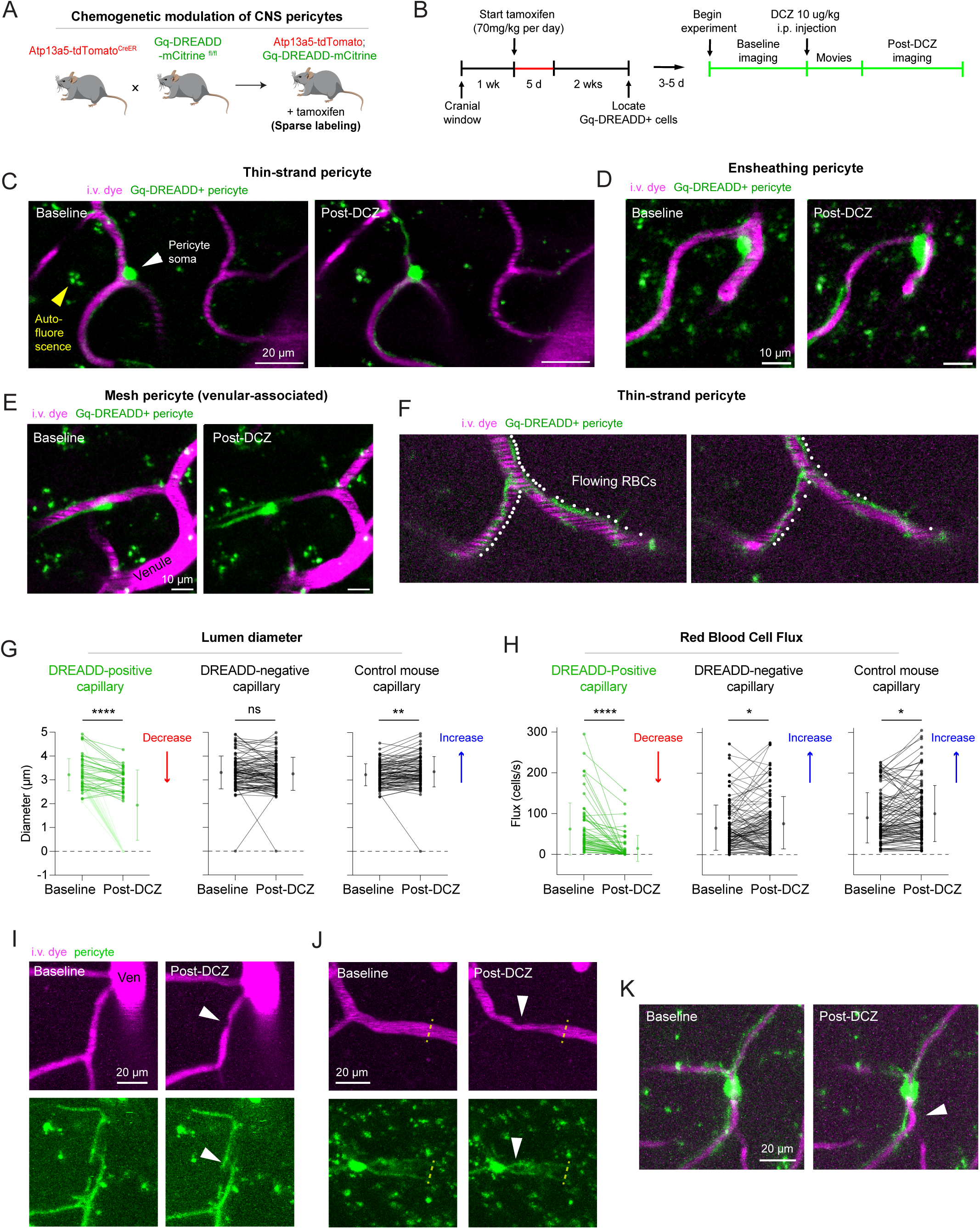
Potent chemogenetic contraction of pericytes in vivo using Gq-DREADD stimulation. **(A)** Cross-breeding to generate mice involved in pericyte chemogenetic experiments. **(B)** Experimental timeline. **(C)** Example of thin-strand pericyte contraction. Gq-DREADD-positive pericyte is green. Non-specific autofluorescence marked by yellow arrowheads. **(D)** Example of ensheathing pericyte contraction. **(E)** Example of contracting mesh pericyte proximal to a venule. **(F)** Reduced number of transiting blood cell shadows following thin-strand pericyte contraction. White dots flank blood cell shadows. **(G,H)** Change in capillary diameter (G) and RBC flux (H) in regions covered by Gq-DREADD-positive pericytes, Gq-DREADD-negative pericytes, and in capillaries of littermate controls. Diameter: **p<0.01, ***p<0.001, linear mixed-effects model, n = 54 capillaries from 8 mice for Gq-DREADD-positive capillary, n = 99 capillaries from 8 mice for Gq-DREADD-negative capillary, n = 96 capillaries from 5 control mice (2 DCZ-treated WT mice, 3 saline-treated WT mice); Flux: *p<0.05, ***p<0.001, linear mixed-effects model, n = 58 capillaries from 8 mice for Gq-DREADD-positive capillary, n = 113 capillaries from 8 mice for Gq-DREADD-negative capillary, n = 127 capillaries from 5 control mice (3 DCZ-treated WT mice, 3 saline-treated WT mice). See Methods for details on statistical approach. **(I)** Secondary processes locally enwrapping the circumference of a peri-venous capillary forms a divot (arrowhead) in the vessel wall during chemogenetic contraction. **(J)** Vasoconstriction does not extend beyond the territory covered by processes of a Gq-DREADD-positive pericyte (dotted line shows end of process, arrowhead shows divot near pericyte soma). **(K)** Longitudinal contraction of thin-strand pericyte process forms a bend in the underlying capillary.

We implanted chronic cranial windows in tamoxifen-treated Atp13a5-Gq-DREADD mice, and several weeks later, *in vivo* two-photon imaging was used to measure capillary diameter and red blood cell (RBC) flux under isoflurane anesthesia (**Fig. 2B**). Imaging was focused on regional capillary networks containing a single Gq-DREADD expressing pericyte (**Fig. 2C-F, Supplementary Fig. 9**). Following intraperitoneal injection of the BBB-permeant DREADD agonist deschloroclozapine (DCZ)^26^, strong capillary constriction and RBC flux reduction was observed specifically in capillary branches contacted by Gq-DREADD+ pericytes (**Fig. 2G,H**). In many cases, there was complete cessation of CBF in constricted capillary segments and the lumen was no longer visible for diameter measurements (reported as zero diameter). Consistent with rapid entry of DCZ into the brain^26^, constrictions initiated within a 2 minutes following systemic DCZ injection and developed over 10-15 minutes (**Supplementary Movies 1-7**). In some cases, pericytes were re-examined ∼1 hour later and constrictions were sustained for at least this period of time. In contrast, nearby capillaries lacking Gq-DREADD+ pericyte coverage exhibited no diameter change, on average, and modest increases in RBC flux (**Fig. 2G,H**). Similar effects were seen across all Gq-DREADD+ mice examined (**Supplementary Fig. 10A,B**). We observed no constrictions in other control groups, which included pooled data from saline vehicle treatment of Gq-DREADD+ mice and DCZ treatment of control littermate mice (**Fig. 2G-H, Supplementary Fig. 10C,D**). A slight vasodilation and increase in RBC flux was seen in capillaries of control mice, consistent with a gradual dilatory effect of isoflurane as shown in our prior studies.^3,27^ These overall data demonstrate the potency and consistency of chemogenetic pericyte contraction as a means to modulate capillary perfusion *in vivo*.

Gq-DREADD+ ensheathing pericytes were occasionally also observed in the ACT zone where, as expected, strong contractility was induced with chemogenetic stimulation (**Fig. 2D, Supplementary Fig. 9**). However, most positive cells were thin-strand pericyte in mid-capillary and peri-venous regions. Remarkably, Gq-DREADD+ pericytes close to ascending venules also exhibited contractility, suggesting that blood flow regulation extends to the distal ends of the capillary bed (**Fig. 2E, Supplementary Fig. 9, Supplementary Movies 1-3**). To better understand the effect of constriction on local blood flow, we measured flow of capillaries upstream and downstream of constricted segments for a qualitative assessment in 8 Atp13a5-Gq-DREADD mice (**Supplementary Fig. 11**). Constrictions in the ACT zone and peri-venous capillaries tended to cause broader regions of hypoperfusion as they gated flow into the capillary bed or drained blood from multiple converging capillaries, respectively. Interestingly, while constrictions in the capillary zone induced local flow loss, they occasionally exhibited blood flow increases in upstream capillaries, possibly due to re-routing of blood cells or compensatory vasodilation in upstream arterioles. Thus, pericyte contraction at the arteriole and venous ends of the capillary bed may have greater effects on capillary perfusion, while compensatory responses may be evoked when mid-capillaries are affected. The magnitude of pericyte contraction engaged by chemogenetic stimulation is supra-physiological, but the results clearly demonstrate the capacity for pericyte-mediated tone generation across the entire capillary bed.

We next asked how pericyte morphology enables vasoconstriction during chemogenetic stimulation. Pericyte contractions were stronger near the pericyte soma since these regions exhibit more pericyte coverage, consistent with prior studies (**Supplementary Fig. 12, Supplementary Movie 1,2,6**).^28^ However, close inspection of thin-strand pericytes revealed that their processes could also distort the vessel despite their partial coverage of the endothelial surface. The processes often twisted around the endothelium, and their tightening during contraction formed “divots” in the lumen, particularly at locations where secondary processes wrapped the lumen (**Fig. 2I, Supplementary Movie 1**). Further, the thin processes could compress the endothelium, leading to reduced lumen diameter (**Supplementary Movie 3-5**). The mechanics of compression could not be resolved, by may be induced by circumferential cinching of thin processes. These diameter changes were specific to the chemogenetically-activated pericyte, as transitions between constricted and non-constricted regions were observed along the same capillary branches where Gq-DREADD+ pericyte processes abutted neighboring non-expressing pericytes (**Fig. 2J, Supplementary Movie 2,3,7**). Tightening and shortening of the processes along the longitudinal axis was also apparent, as distortion of the capillary lumen could be seen when processes pulled toward the soma (**Fig. 2K, Supplementary Movie 1,3,4-7**). These data reveal how thin pericyte processes use a variety of structural mechanics to alter endothelial shape, thereby affecting lumen size and blood flow.

### Sparse pericyte contraction results in distributed hypoxic microdomains

We next used Hypoxyprobe immunohistology to determine whether sparse pericyte contraction resulted in regional hypoxia. Hypoxyprobe binds tissues with pO_2_ < 10 mm Hg following injection *in vivo*, and labeled hypoxic tissues can be detected post-mortem with immunohistology.^29^ In saline-treated control mice, Hypoxyprobe labeling was low and uniform in most brain regions examined (**Fig. 3A-E, left panels**). Higher basal labeling was occasionally seen in the hippocampus, particularly in pyramidal neuron layers, possibly due to the combination of high metabolic activity and lower vascular density.^30^ However, in DCZ-treated mice, scattered microdomains of hypoxia were apparent throughout the brain (**Fig. 3A-E, right panels**), likely reflecting domains of tissue affected by the contraction of isolated pericytes. Numerous limbic system structures exhibited higher densities of hypoxic microdomains, including the hippocampus, lateral cortical regions (including temporal association, entorhinal, peri-rhinal), basolateral brain regions (including striatum-like amygdalar nuclei, basolateral amygdalar nuclei within the cortical subplate, and piriform cortex)(**Fig. 3F-H; Supplementary Fig. 13A-E**). These data reveal that limbic structures are vulnerable to capillary perfusion deficits, while other regions, such as the more highly vascularized somatosensory cortex (S1) and thalamus (**Fig. 3I,J**), are more resilient.

**Figure 3.**
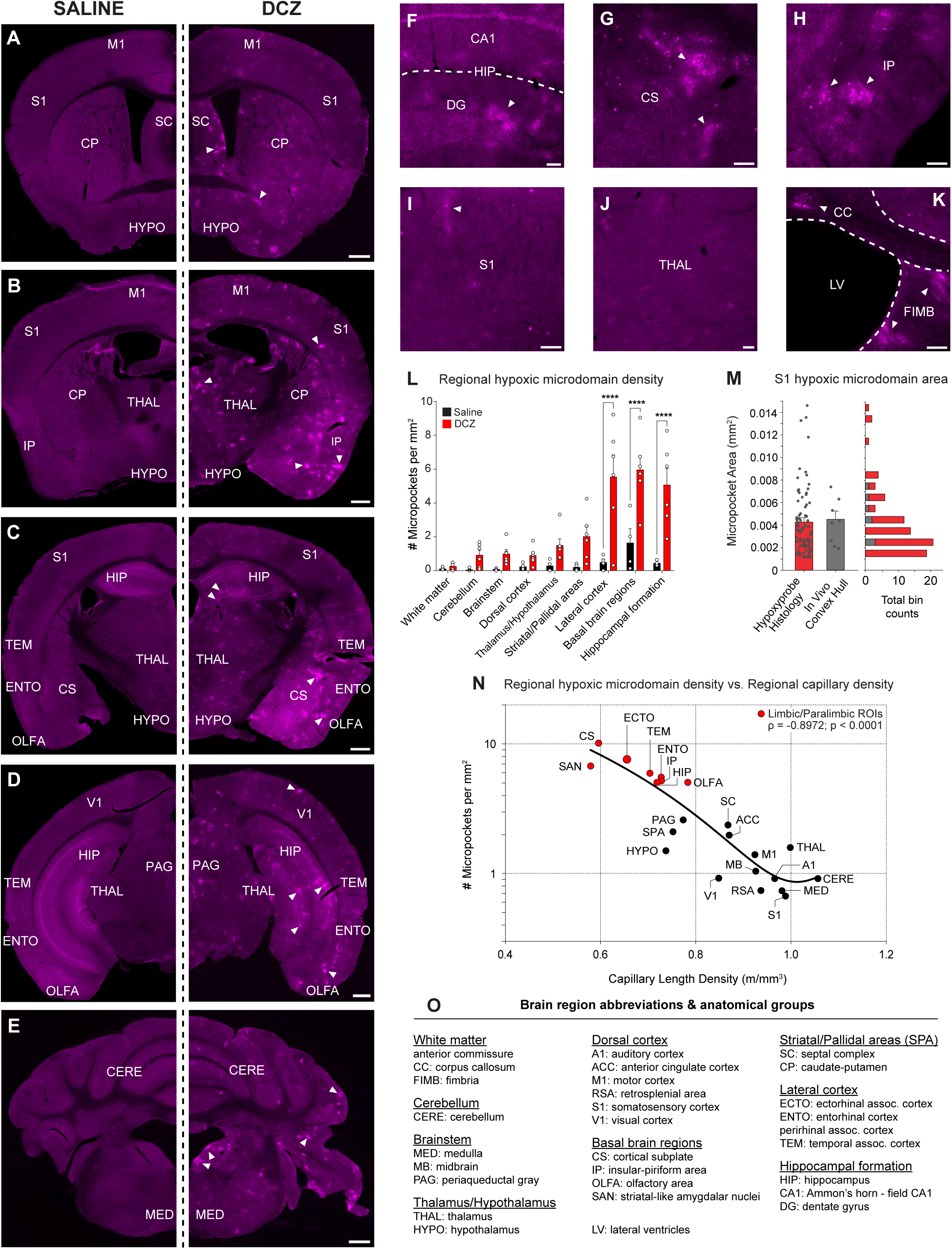
Post-mortem characterization of regional tissue hypoxia after sparse chemogenetic contraction of CNS pericytes. **(A-E)** Hypoxyprobe-labeled coronal sections from saline control-treated (left) and DCZ-treated (right) Atp13a5-Gq-DREADD brains along the anteroposterior axis. White arrows show examples or discrete microdomains of tissue hypoxia. All scale bars: 500□μm. **(F-K)** Hypoxyprobe staining in DCZ-treated Atp13a5-Gq-DREADD mice, showing CA1 and dentate gyrus of the hippocampus (F), cortical subplate (G), insular-piriform area (H), somatosensory cortex (I), thalamus (J), and corpus callosum and fimbria white matter tracts (K). All scale bars: 100 μm. **(L)** Comparison of regional hypoxic microdomain density across nine anatomical groups (defined in panel O) in DCZ-treated (n = 6 mice, red bars) vs saline-controls (n = 4 mice, black bars) mice using two-way ANOVA with *post-hoc* comparisons. Treatment main effect: f(1,72) = 56.31, ****p < 0.0001; Anatomical group main effect: f(8,72) = 9.416, ****p < 0.0001; Treatment and anatomical group interaction: f(8,72) = 4.92, ****p < 0.0001. *Post-hoc* comparison (Šídák correction) of hypoxic microdomain density between DCZ-treated and saline-controls across anatomical groups: lateral cortex, basal brain regions, hippocampal formation (all ****p < 0.0001). All other groups: n.s, p > 0.05. See Methods for details on statistical approach. Bar graph data presented as mean ± SEM. **(M)** Comparison of hypoxic microdomain density identified with Hypoxyprobe histology (red bars) and convex hull area (gray bars) representing regional blood flow cessation captured by *in vivo* two-photon imaging in the somatosensory cortex (S1) after DCZ treatment (left panel). Hypoxyprobe histology (n = 87 measurements from 6 mice; mean ± SEM: 0.0043 ± 3.79 x 10^-4^ mm^2^); *in vivo* convex hull (n = 8 Gq-DREADD-positive regions from 8 mice; mean ± SEM: 0.0045 ± 7.39 x 10^-4^ mm^2^). Frequency counts binned by microdomain area size are plotted as the proportion of all total measurements (right panel). P = 0.3, Mann Whitney test. Bar graph data presented as mean ± SEM. **(N)** Regional hypoxic microdomain density (this study) plotted as a function of capillary length density measurements from Ji et al.^2^ Data was fitted with a second order decay quadratic model, which demonstrates a strong inverse relationship (see Methods for details). Goodness of fit: R^2^ = 0.798, adjusted R^2^ = 0.776. Spearman’s ranked correlation test (non-parametric): Spearman’s ρ = -0.897, 95% CI [-0.958, -0.759], ****p < 0.0001 (two-tailed). **(O)** Legend of brain region abbreviations and anatomical groups.

Basal Hypoxyprobe labeling was distinctly lower in white matter, likely due to reduced probe binding affinity. However, hypoxic microdomains were still visible in the corpus callosum, fimbria, and other major fiber tracts in DCZ-treated mice, though less frequent than in gray matter (**Fig. 3K, Supplementary Fig. 13F-I**). This indicates that white matter pericytes can also contract. Hypoxic microdomains were also observed within the periaqueductal grey of the midbrain, which receives afferents from the amygdala (**Supplementary Fig. 13J**). Interestingly, enriched clusters of hypoxic tissue were found in the medial vestibular nucleus lining the 4th ventricle in the cerebellum, which is critical for balance and posture (**Supplementary Fig. 13K**). Labeling also appeared to be enriched in the lateral septal nucleus, particularly the caudo-dorsal region lining the lateral ventricle (**Supplementary Fig. 13L**), which forms connections to the hippocampus.

To understand whether hypoxic microdomains resulted from contraction of isolated pericytes, we next quantified the size of individual hypoxic microdomains in the somatosensory cortex, comparing areal measurements from Hypoxyprobe histology (**Fig. 3M, red**) to measurements approximating the area of blood flow loss produced by pericyte contraction *in vivo* (**Fig. 3M, gray**). For the latter analysis, a convex hull area was bounded by distal branch-points of capillary segments experiencing ≥ 50% blood flow reduction from baseline. The average microdomain size was remarkably similar between histology (0.0043 ± 3.79 x 10^-4^ mm^2^) and *in vivo* measurements (0.0045 ± 7.40 x 10^-4^ mm^2^), suggesting that hypoxic microdomains detected histologically likely resulted from contraction of individual pericytes.

### Hypoxic burden is strongly related to regional microvascular density

Regional variation in tissue hypoxia may reflect differences in capillary density. Sparser capillary networks offer less redundancy to circumvent focal obstructions, leading to increased likelihood of hypoxia. To investigate this possibility, hypoxic microdomain density was plotted as a function of regional capillary density measurements, reported recently in a mouse brain vascular connectome (**Fig. 3N**).^2^ This prior work from Ji *et al.* revealed that neocortical regions are highly vascularized, whereas basolateral regions including the amygdala, cortical subplate, and paleocortex (including piriform cortex) contain ∼30% lower average vessel density. In our analysis, hypoxic microdomain density was strongly inversely correlated with regional capillary length density across surveyed brain regions. Notably, multiple limbic system regions exhibited low vascular density and accordingly the highest levels of Hypoxyprobe labeling (**Fig. 3N**).

We also measured the proportion of total Gq-DREADD+ pericytes across regions of high and low capillary density (**Supplementary Fig. 8C**). Gq-DREADD+ pericyte density was similar between regions of higher and low hypoxic burden. A marginally higher recombination rate was detected in the cortical subplate (low capillary density) compared to the thalamus (high capillary density). However, the magnitude of difference was small, at ∼3% vs 6% of total pericytes in thalamus vs. cortical subplate, respectively. Thus, we conclude that regional capillary density is the primary determinant of hypoxic outcome.

### Capillary flow defects elude detection by conventional MRI perfusion imaging

To understand how sparse pericyte contraction affects CBF on a broader level, we performed arterial-spin labeling MRI. Isoflurane-anesthetized Atp13a5-Gq-DREADD mice (and non-expressing controls) were imaged under baseline conditions and 10-minutes following intraperitoneal DCZ injection (**Fig. 4A**). Consistent with known differences in vascular density, there was a significant effect of region on baseline CBF. The amygdala and hippocampus exhibited significantly lower baseline CBF than regions with higher microvascular density, including the thalamus and hypothalamus (**Fig. 4B**). However, we found no difference in DCZ-induced CBF change between experimental groups on a brain-wide or regional level. Overall, these data suggest that regional capillary flow deficits can be masked by broader CBF signals when using lower-resolution imaging techniques such as MRI.

**Figure 4.**
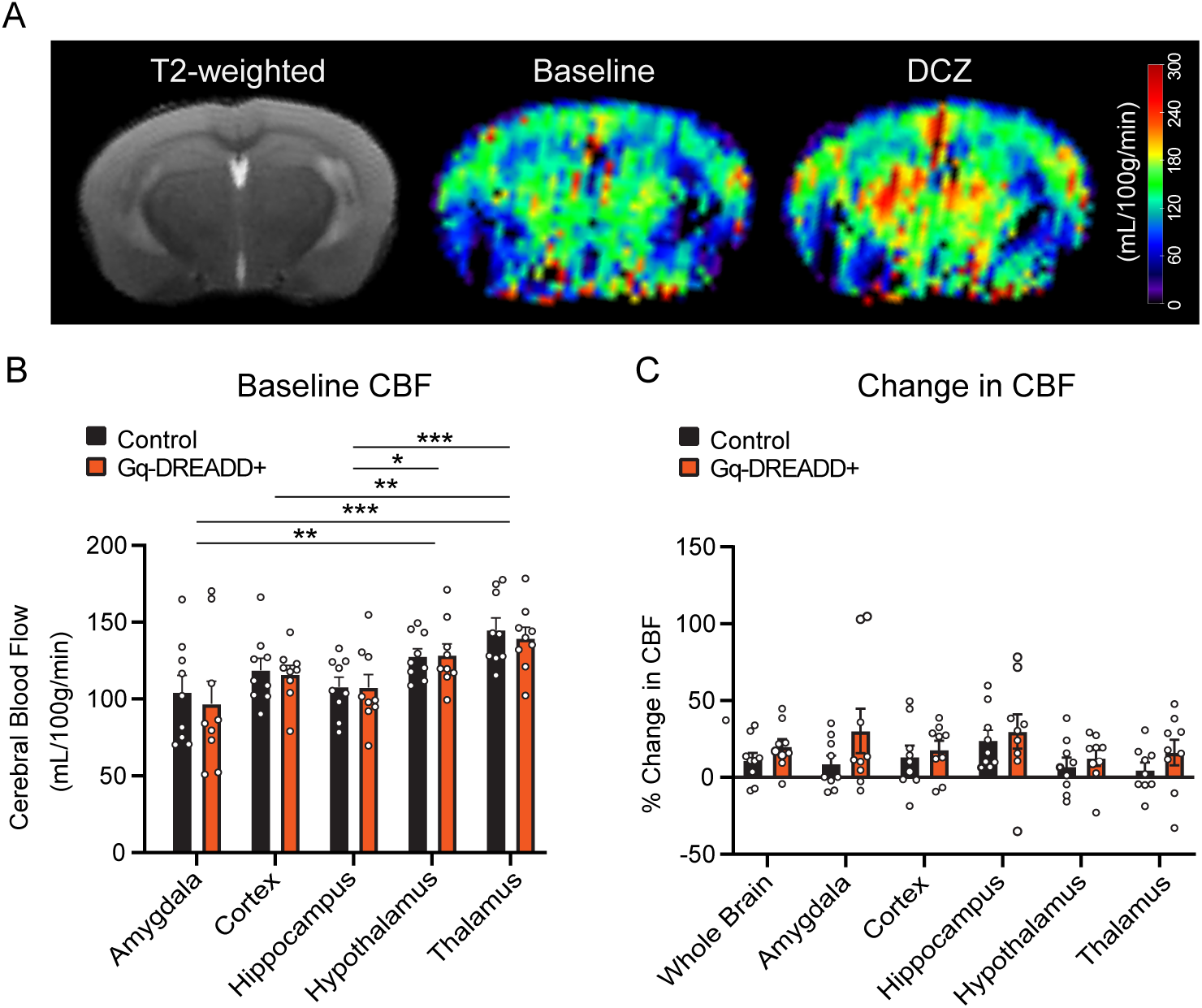
MRI-based CBF measurements in mice with sparse pericyte contraction. **(A)** Structural and CBF imaging by MRI. 7T T2-weighted image showing brain anatomy (left panel). Representative ASL images of CBF from Atp13a5-Gq-DREADD mouse at baseline (middle panel) and 15-30 min post-DCZ administration (right panel). **(B)** Regional CBF levels at baseline in control WT mice and Atp13a5-Gq-DREADD mice. A linear mixed-effects model revealed a significant main effect of Region, F(5, 80) = 14.35, p < .001. There was no main effect of Group, F(1, 16) = 0.11, p = 0.75, and no Region × Group interaction, F(5, 80) = 0.16, p = 0.98. N = 9 mice per group. For *post-hoc* analysis *p<0.05, **p<0.01, ***p<0.001. Data show pairwise comparison outcomes, collapsed across groups. See Methods for details on statistical approach. **(C)** Change in CBF over baseline observed at 15-30 min after DCZ administration. There were no significant effects of Region, F(5, 75.70) = 0.93, p = 0.47, Group, F(1, 9.76) = 2.16, p = 0.17, or Region × Group interaction, F(5, 70.95) = 0.43, p = 0.82. N = 9 mice per group. See Methods for details on statistical approach.

## DISCUSSION

In this study, we leveraged sparse Gq-DREADD expression in CNS pericytes to understand their contractile capacities and effect on tissue oxygen across the brain. We find that pericytes across the capillary bed can influence lumen diameter and blood flow, including pericytes near venules. Capillary pericytes can tighten circumferentially at cell bodies and pull their processes longitudinally to exert force on the endothelium, and their contraction is sufficient to generate regional hypoperfusion and microdomains of tissue hypoxia. These hypoxic microdomains are enriched in multiple brain regions related to limbic system function, including hippocampus, amygdala, piriform cortex, lateral septal nucleus, and the vulnerability of these regions is linked to lower basal microvascular densities in these regions. This suggests that limbic structures are less resilient to capillary flow defects and are more prone to poor blood flow and oxygenation when challenged.

Our characterizations of the Atp13a5-CreERT2 mouse line support its use for genetic targeting of CNS capillary pericytes^21,23^, including mesh and thin-strand subtypes with low or undetectable α-SMA expression, as well as a subset of ensheathing pericytes, which are α-SMA-positive. Using the strong YFP reporter line Ai3, we observed no labeling in smooth muscle cells of cerebral arteries or arterioles, or in microvascular beds of peripheral organs examined. Standard tamoxifen dosing labeled ∼50–60% of brain pericytes, with lower targeting in spinal cord and retina. In contrast, recombination efficiency in the floxed Gq-DREADD line was only 1–9%, likely reflecting non-ROSA26 transgene insertion. Consequently, genetic manipulations will yield mosaic targeting and require careful characterization. Despite this limitation, the Atp13a5-CreERT2 line enables more precise genetic dissection of CNS pericyte function than other mural cell targeting mouse lines.

CNS pericytes express a diverse repertoire of GPCRs.^24^ Prior brain slice studies show that Gq-coupled agonists, including endothelin-1 and the thromboxane receptor agonist U46619, induce pericyte contraction.^11,31–33^ Our *in vivo* Gq- studies demonstrate that Gq signaling alone is sufficient to drive potent pericyte-mediated capillary constriction, indicating that contractility is an intrinsic feature of this pathway rather than receptor-specific. The physiological and pathophysiological roles of individual GPCRs in pericyte function remain to be defined. Under basal conditions, tonic Gq-GPCR activity may help maintain steady-state capillary tone and proper blood flow distribution.^34–36^ In disease, aberrant Gq signaling may promote vasoconstriction and flow resistance, similar to the supra-physiological stimulations performed here^11^ Resolving these mechanisms will require more CNS pericyte-specific knockout studies, as prior work has largely relied on less selective pharmacological manipulations.

A longstanding question in the blood flow field is how thin-strand pericytes generate contractile force despite their fine processes and undetectable expression of α-SMA.^37^ Our *in vivo* imaging reveals that pericytes can exert substantial force at both their somata and processes, at times fully collapsing the capillary lumen. Although optical resolution remains limiting, constriction may occur through circumferential “cinching” by thin processes. We also observed longitudinal contraction, with process tightening creating divot-like resistance points and bending of the capillary. Thus, pericyte cytoskeletal networks, even with relatively low actomyosin content, can generate sufficient force given the low intraluminal pressure and compliant endothelial layer. Comparable mechanisms have been described in retinal capillary pericytes, and brain pericytes appear to operate similarly.^38–40^

In a recent study, Wu et al. used the same mouse cross (Atp13a5-CreERT2-IRES-tdTomato × floxed Gq-DREADD) and reported a paradoxical increase in cerebral blood flow following acute chemogenetic stimulation.^24^ Although this appears inconsistent with our results, and with the established view that Gq-GPCR signaling drives mural cell contraction, we suspect the discrepancy arises because the locations of Gq-DREADD+ pericytes were not determined during imaging. Without this information, blood flow responses to sparse pericyte contraction could reflect compensatory changes elsewhere in the network, similar to those observed in neighboring control vessels in our dataset.

In humans, limbic system degeneration occur with aging and is exacerbated in cerebral small vessel disease and Alzheimer’s disease.^41–43^ Our findings raise the possibility that intrinsic limitations of microvascular architecture contribute to the early vulnerability of brain regions essential for cognition, emotion, memory, and social behavior. Increased gliosis in the amygdala in small vessel disease supports a vascular contribution to regional pathology.^44^ Studies in aging mice further report selective pericyte loss in basal forebrain and amygdalar regions, consistent with localized vascular deficits.^20^ Lower microvascular density in limbic structures limits alternative flow pathways during capillary obstruction by pericytes or circulating leukocytes.^6^ These regions may also have fewer ACT and venous branches to support blood flow input and output, increasing susceptibility to perfusion bottlenecks. The microvascular organization of limbic structures remains incompletely characterized in both humans and rodents and warrants further investigation.

## ACKNOWLEDGMENTS

Projects in the Shih lab were supported by grants from the NIH/NIA (R01AG062738, R01AG077731, R01AG081840, R61NS137365) and the Leducq foundation (23CVD03). DTC was supported by an American Heart Association Pre-doctoral fellowship (26PRE1560770). SKB was supported by fellowships from the NIH/NINDS (F32NS117649) and NIH/NIA (K99AG080034). DD was supported by NIH/NINDS grant R01NS112526. LF and JG were supported by grants from the NIH/NINDS (K08NS118138) and NIH/NCI (R37CA275954). We thank David Kleinfeld, Yan Wu, and Meng Lui for valuable discussions.

## AUTHOR CONTRIBUTIONS

LTS and AYS conceptualized the study. Experiments and data analysis were performed by LTS, DTC, CF, BZ, KE and LF. Manuscript was written by AYS with editing and contributions from all authors.

## STAR METHODS

## Key Resources Table

| Reagent or Resource | Source | Identifier |
| --- | --- | --- |
| **Chemicals, antibodies, and reagents** |  |  |
| Alexa fluor 680 carboxylic acid, succinimidyl ester | Thermo Fisher Scientific | A20008 |
| Anti-α-Smooth Muscle antibody, Alexa fluor 647 conjugated | R & D Systems | IC1420R-100UG |
| Anti-HA-tag antibody, Rabbit monoclonal | Cell Signaling Technology | 3724T |
| Anti-PDGFRβ, Rat monoclonal | Thermo Fisher Scientific | 14-1402-82 |
| Anti-rabbit Alexa fluor 488 antibody (goat host) | Thermo Fisher Scientific | A11008 |
| Anti-rabbit Alexa fluor 647 antibody (goat host) | Thermo Fisher Scientific | A21244 |
| Anti-rabbit Alexa fluor 647 (donkey host) | Abcam | Ab150075 |
| Anti-rat Alexa fluor 647 (goat host) | Thermo Fisher Scientific | A21247 |
| DAPI | Sigma-Aldrich | D9542 |
| Deschloroclozapine | Hello bio | HB9126 |
| Fluoromount-G | Thermo Fisher Scientific | 00-4958-02 |
| Hypoxyprobe Omni Kit (1000 mg pimonidazole HCl plus 2 units of pab2627 rabbit antisera) | Hypoxyprobe | HP3-1000Kit |
| Normal goat serum | Sigma-Aldrich | G9023 |
| Nycodenz | Fisher Scientific | 100334-594 |
| OCT compound | Fisher Scientific | 23-730-571 |
| Ophthalmic ointment | Amazon | B0CGY3XXFT |
| Paraformaldehyde (4%) | Thermo Fisher Scientific | J19943.K2 |
| Phosphate buffered saline | Sigma-Aldrich | P3813 |
| Sodium azide | Sigma-Aldrich | S2002 |
| Sucrose | Sigma-Aldrich | S7903 |
| Tetrahydrofuran | Sigma-Aldrich | 186562 |
| TrueBlack | Biotium | 23007 |
| Urea | Sigma-Aldrich | U5378 |
| **Experimental models: organisms/strains** |  |  |
| Atp13a5-CreERT2-IRES-tdTomato | Zhen Zhao lab (USC) | N/A |
| Ai3 mice | The Jackson Laboratory | Strain #: 007903 |
| CAG-LSL-Gq-DREADD mice | The Jackson Laboratory | Strain #: 026220 |
| **Surgical consumables** |  |  |
| Atipamezole (antisedan) | Patterson Veterinary | 07-867-7097 |
| Betadine | Fisher Scientific | NC1696484 |
| Buprenorphine | Patterson Veterinary | 07-894-9214 |
| Burrs for micro drill | Fine Science Tools | 19007-05 |
| Cyanoacrylate glue (Loctite 401) | Uline | S-15623 |
| Dexmedetomidine hydrochloride (dexdormitor) | Patterson Veterinary | 07-867-7105 |
| Ethanol | Decon labs | V1001 |
| Fentanyl citrate | Patterson Veterinary | 07-890-5657 |
| Flumazenil | Patterson Veterinary | 07-892-5077 |
| Meloxicam | Patterson Veterinary | 07-891-7959 |
| Metabond Dental Cement (C&B-Metabond®) | Patterson Dental | 75533484 |
| Midazolam | Patterson Veterinary | 07-894-9225 |
| Optical glue, UV-curable | Edmund Optics | 37-322 |
| Round coverslip (3 mm) | Warner Instruments | 64-0720 |
| Round coverslip (4 mm) | Warner Instruments | 64-0724 |
| **Software and algorithms** |  |  |
| Advanced Neuroimaging Tools (ANTS) | Avants et al.<sup>45</sup> | N/A |
| FMRIB Software Library (FSL) | Jenkinson et al.<sup>46</sup> | N/A |
| Illustrator 2025 Version 29.0.1 | Adobe | N/A |
| ImageJ, Fiji Version 2.1.0/1.53c | https://imagej.net/ij/ | N/A |
| IMARIS | Oxford Instruments | N/A |
| MATLAB_R2025a Version 25.1.0.2943329 | MathWorks | N/A |
| Prism, Version 10.0.3 | GraphPad Software | N/A |
| R (ver 4.5.2) | https://www.r-project.org/ | N/A |

### Animals

Mice were housed in specific-pathogen-free facilities approved by AALAC and were handled in accordance with protocols approved by the Seattle Children’s Research Institute and Oregon Health and Sciences University IACUC committees. Mice were housed in plastic cages on a 12 h light/dark cycle with access to water ad libitum and a standard laboratory diet.

For YFP reporter studies involving brain region analyses, we generated Atp13a5-CreERT2:Ai3 mice by cross-breeding Atp13a5-CreERT2-IRES-tdTomato mice^21^, originating from the lab of Zhen Zhao, with Ai3 (floxed YFP reporter) mice from Jacksons Lab (Strain #: 026220, C57BL/6 background)^47^. Six bigenic mice (3-4 month old females) were used to analyze endogenous YFP expression in Atp13a5 positive cells across the CNS and peripheral organs. Three mice were used to define genetic targeting in the capillary and ACT zones. For both studies, mice were treated with 70 mg/kg of tamoxifen, dissolved in corn oil, daily via intraperitoneal (I.P.) injection for five consecutive days. Approximately one month post tamoxifen dosing, mice were trans-cardially perfused with PBS and 4% PFA under deep anesthesia (Euthasol). For profiling across the CNS and peripheral organs, brain, eyes, spinal cord, skull, heart, lung, liver, kidney, skeletal muscle (gastrocnemius), stomach and intestine (duodenum) were collected and placed in PBS. All organs except the eyes, were post-fixed in 4% PFA for 90 minutes and transferred to 30% sucrose in PBS for 48 hours for cryopreservation. Whole retina was dissected from eyes in PBS and directly slide mounted. All other organs were cryosectioned coronally at 40 µm thickness and directly slide mounted. To characterize genetic tarting of the ACT zone, brains were collected, post-fixed in 4% PFA overnight and then sectioned into 1-2 mm slabs using a mouse brain matrix. They were then processed for tissue clearing and imaging, as described below.

For chemogenetic studies, we generated Atp13a5-Gq-DREADD mice by cross-breeding Atp13a5-CreERT2-IRES-tdTomato mice with CAG-LSL-Gq-DREADD mice (floxed Gq-DREADD) mice from Jacksons Lab (Strain #: 026220, C57BL/6 background).^48^ Eight mice were used for *in vivo* two-photon imaging. Controls included 3 saline-treated Gq-DREADD-positive mice and 3 DCZ-treated wild-type littermates, which were combined into a single control group. Ten Atp13a5-Gq-DREADD mice were used for *post-mortem* Hypoxyprobe immunohistology, with 6 receiving DCZ and 4 receiving saline control.

### Cranial window surgery

For chronic cranial window implantation, dexamethasone (2□μg per g of body weight; Patterson Veterinary) was subcutaneously administered to animals 4□h before the surgery, which helped to reduce brain swelling during the craniotomy. Anesthesia was induced with a cocktail consisting of fentanyl citrate (0.05 mg/kg), midazolam (5□mg/kg) and dexmedetomidine hydrochloride (dexdormitor, 0.5□mg/kg), and buprenorphine was given pre-operatively for analgesia (0.1 mg/kg). The scalp was cleaned with betadine and ethanol. Then under sterile conditions, the scalp was excised and the periosteum cleaned from the skull surface. An aluminum flange for head fixation was attached to the right side of the skull surface using Loctite 401 cyanoacrylate glue and C&B-Metabond quick adhesive cement. A circular craniotomy (dura left intact), ∼3□mm in diameter, was created over the left hemisphere and centered over 2□mm posterior and 3□mm lateral to bregma. The craniotomy was sealed with a glass coverslip plug consisting of a round 3-mm glass coverslip glued to a round 4-mm coverslip with ultraviolet light-cured optical glue. The coverslip was positioned with the 3-mm side placed directly over the craniotomy, whereas the 4-mm coverslip was laid on the skull surface at the edges of the craniotomy. Loctite 401 was carefully dispensed along the edge of the 4-mm coverslip to secure it to the skull. The area around the cranial window was then sealed with dental cement. Throughout the surgery, the body temperature was maintained at 37□°C with a feedback-regulated heat pad (FHC Inc.) and mice were provided medical air through a nose cone (20–22% oxygen and 78% nitrogen, moisturized by bubbling through water; AirGas Inc.). The anesthesia was reversed post-surgery by s.c. injection of a mixture containing atipamezole (antisedan, 2.5 mg/kg), flumazenil (0.5 mg/kg), and buprenorphine (0.1 mg/kg) diluted in sterile Lactated ringers (1 mL). Mice were allowed to regain consciousness before returning to the vivarium. Imaging was initiated after a 3-week recovery period.

### Two-photon microscopy

*In vivo* two-photon imaging system was performed with a Bruker Investigator (run by Prairie View 5.5 software) coupled to a Spectra-Physics Insight X3. Green (YFP or mCitrine), red (tdTomato), and far-red (Alexa-680-dextran) fluorescence emission was collected through 525/70 nm, 595/50 nm and 660/40 nm bandpass filters, respectively, and detected by GaAsP photomultiplier tubes (PMTs). High-resolution imaging was performed using a water immersion 20-X, 1.0 NA objective lens (Olympus XLUMPLFLN 20XW) using 975 nm excitation during the experiment, with laser powers maintained below 40 mW. Lateral sampling resolution was 0.4 μm per pixel and axial sampling was 1μm steps between frames. To label the vasculature, a custom-conjugated Alexa Fluor 680-dextran (Alexa Fluor 680; Life Technologies; A20008 conjugated to 2 MDa Dextran; Fisher Scientific; NC1275021)^27^ was injected through the retro-orbital vein under deep isoflurane anesthesia. During imaging, isoflurane was maintained at 1.5% MAC in medical-grade air.

### Imaging timeline and chemogenetic stimulation

Two-photon imaging, as described above, was used to obtain baseline metrics and to study the effects of pericyte chemogenetic activation. Pre-imaging was performed to identify locations of Gq-DREADD-positive pericytes (based on Gq-DREADD-mCitrine fusion protein detection) 3-weeks after cranial window implantation and 2-7 days prior to experiments involving chemogenetic stimulation. During chemogenetic experiments, Gq-DREADD-positive pericytes identified from pre-imaging were re-located and Z-stacks were collected, centered around these pericytes. Following z-stacks, linescans were obtained along the centerline of as many capillaries as achievable within the region of interest over a 15-20 minute period. These line-scans were collected from capillaries contacted by the Gq-DREADD-positive pericyte and surrounding capillaries within the same local network. Line-scans were later used for assessment of RBC flux at baseline. Deschloroclozapine was then injected intraperitoneally at a dose of 10 µg/kg in PBS (0.1 mL volume). Injections were performed gently as the mice remained under the microscope. Immediately thereafter (within 1-2 minutes of IP injection), successive image stacks were taken over a 10-15 min window to capture dynamic changes in pericyte contractile state. After completion of time-series collection, post-DCZ Z-stacks and capillary line-scans were captured in the same locations as during baseline imaging.

### Analysis of in vivo vascular metrics

The diameters of different vessel types were measured in maximally projected images from high-resolution Z-stacks. Maximum projections of 10-40 µm in thickness were used for analysis, dependent upon the size of the vessels. To reduce bias of measurement location, we used a custom ImageJ/Fiji macro called VasoMetrics to analyze lumen full-width at half maximum diameter at multiple, equidistant locations (spaced 1 µm) along each vessel segment of interest.^49^ The values across each vessel segment were used to calculate the average diameter of the vessel and the standard deviation of diameter along the measured region. Line-scanning was performed at 3-X digital zoom to guide accurate placement of the scan. Vessel segments were sampled individually with line-scan duration set to ∼1.2 s at a sampling frequency of ∼2 kHz. For each line-scan captured, we calculated RBC flux by manually counting the number of blood cell shadows over the total duration of the line scan and represent the values as cells per second. For convex hull analysis of area of decreased blood flow, we determined all capillary segments within a network that experienced ≥ 50% blood flow reduction from baseline. In maximally-projected images, the distal ends of these capillary segments were used as points to demarcate a polygon, and the area enclosed by the polygon was reported as an estimated hypoxic microdomain area.

#### Statistics for In vivo two-photon imaging data

Capillary diameter and blood cell flux measurements were analyzed using a linear mixed-effects model to account for the repeated-measures and nested structure of the data. The fixed effect was stimulus condition (pre- vs. post-DCZ), and random intercepts were included for both mouse ID and capillary ID nested within mouse to account for inter-mouse and inter-capillary variability. For analyses of the effect of the stimulus, capillary diameter was modeled as a function of stimulus condition:

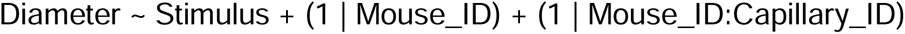

Models were fitted using the lmer function from the lmerTest package in R, which provides approximate p-values for fixed effects. Statistical significance was defined as p < 0.05. Analyses were conducted in RStudio (version 2025.09.2+418).

### PDGFRβ immunostaining

To visualize pericytes, sections from the CNS and peripheral organs were incubated in rat anti-PDGFRβ antibody (1:200; Thermo Fisher Scientific, 14-1402-82), diluted in antibody buffer (1% w/v Triton-X-100, 1% w/v goat serum, 0.01% w/v sodium azide and PBS), for 48 hours at 4°C. Sections were then washed with PBS three times and incubated in goat anti-rat Alexa fluor 647nm (1:1000; Thermo Fisher Scientific, #A21247), diluted in PBS, for 2 hours at room temperature (RT). Sections were washed in PBS and incubated in 0.2% TrueBlack in 70% ethanol for 15 min at RT to quench autofluorescence. Sections were washed in PBS twice and then mounted in Fluromount G containing DAPI, cover slipped and left to dry. Sections were imaged with an Olympus APX100 slide scanner at 20-X magnification at a z-range of 20 µm across five focal planes, spaced 3 µm apart. Optimal exposure times for DAPI (excitation [Ex]: 388 nm; emission [Em]: 448 nm), tdTomato (Ex: 576 nm; Em: 625 nm), YFP (Ex: 494 nm; Em: 530 nm) and PDGFRβ (Ex: 640 nm; Em: 690 nm) were determined manually and kept consistent across all images. *Analysis.* Image post-analysis was performed using Qupath 4.0. Atp13a5-tdTomato positive cells that co-localized with PDGFRβ immunostaining and YFP positive cells that co-localized with Atp13a5-tdTomato, in different regions of the brain, spinal cord and retina, were manually counted to determine cell number per mm^2^. For PDGFRβ positive cells, only cells residing on capillaries were counted to prevent counting of PDGFRβ-positive smooth muscle cells or fibroblasts residing on arterioles. Several regions of interest were placed in the olfactory bulb, striatum, sensory cortex, corpus callosum, hippocampus, thalamus, cerebellum and brain stem of both hemispheres of the brain, in the gray and white matter of the spinal cord, and the retina. Manual counts were averaged for each region per animal.

### HA-tag immunostaining and cell quantification

*Brain slice processing.* A total of n=7 Atp13a5-Gq-DREADD mice and n=3 littermate controls were injected with tamoxifen at 70 mg/kg daily for 5 consecutive days and allowed to express transgene for several weeks before trans-cardial perfusion and tissue collection. Mice were anesthetized with 5% isoflurane and maintained at 2% isoflurane during 5-minute transcardial perfusion with room temperature (RT) 0.9% saline containing 10 mg/mL heparin followed by a 5-minute perfusion with RT 4% paraformaldehyde (PFA). Extracted whole brains were fixed overnight in 4% PFA at 4°C and stored in 1x PBS before being cut into 50 μm sections using a Vibratome (Leica VT 1000S) and stored at 4°C in 0.01% w/v sodium azide in 1x PBS. *Immunostaining*. Three sections corresponding to images 69-73 of the Allen Mouse Brain Atlas (atlas.brain-map.org) from each Atp13a5-Gq-DREADD mouse underwent a standard immunofluorescence protocol. Briefly, sections were washed in 1x PBS and permeabilized in 0.1% Triton-X in 1x PBS (1x PBST) for 10 min before blocking with 10% normal goat serum in PBST for 1 hour with shaking at RT. Sections were then incubated overnight with shaking at RT in the primary antibody (Rabbit anti-HA-tag, Cell Signaling Technology Cat: 3724T) diluted 1:500 in antibody solution (2% Triton-X, 10% normal goat serum in 1x PBS). The next day, sections were washed in 0.1% PBST for 3 x 10 minutes before incubating overnight with shaking at RT in the secondary antibody (Donkey anti-Rabbit Alexa fluor 647, Abcam Cat: ab150075) diluted 1:1000 in antibody solution. The following day, sections were washed in 0.1% PBST for 3 x 10 minutes followed by a 5-minute incubation in 1:1000 DAPI in 1x PBS. Sections were then mounted using Fluoromount-G and left at RT overnight to dry. *Imaging.* Images were captured by a BC43 benchtop spinning disc confocal microscope (Oxford Instruments). The Allen Atlas was consulted to identify brain regions of interest in 3-4 hemispheres per animal. On each hemisphere, 2-3 non-overlapping, 621 µm x 621 µm images were captured within the boundaries of each region at 20-X magnification using identical imaging parameters. *Image Analysis*. Based on background fluorescence, as determined by secondary only antibody labeling, the far-red channel of all images underwent a background subtraction step. The Spots asset of IMARIS 10.2.0 was then used to identify tdTomato+ cell bodies and classify them as HA-tag+ or HA-tag-according to mean Alexa Fluor 647 intensity. Classification parameters for each imaging batch were determined using “training” images taken during each imaging session. A training image was captured on each hemisphere, and these were not included in the final analysis. Ideal parameters were found manually for each training image, and averages used to analyze the corresponding batch. Following an audit of the output, total tdTomato+ cell counts were normally distributed (Shapiro-Wilk test, p=0.2839). Images that undercounted tdTomato+ cells by >25% (>2 standard deviations from the mean) and those that overcounted by >10% (>2.5 standard deviations from the mean) were excluded from analysis. Gq-DREADD recombination rate for each image was calculated as the percentage of tdTomato+ cells that were also HA-tag+ (% HA+). Summary data was obtained by averaging % HA+ cells across animals for genotype effect, or across region per animal for region effect. Genotype data were analyzed using an unpaired t-test with Welch’s correction. Regional data were analyzed using a mixed effects analysis with Geisser-Greenhouse correction, followed by Tukey’s multiple comparisons test. Statistics performed with GraphPad Prism 10.3.1.

### α-SMA immunostaining and EZ-Clear Protocol

To better delineate the boundaries of α-SMA expression within the vascular network, 1-2 mm thick coronal brain tissue slabs from Atp13a5-CreERT2-tdTomato: x Ai3-YFP^fl/fl^ mice (n = 3) were immunostained with an anti-α-SMA antibody with Alexa fluor 647 conjugate, as described previously, with modifications^50^, to perform three channel imaging of tdTomato, YFP and α-SMA. Immunostained tissue slabs were then optically cleared using the EZ-Clear protocol.^51^ Briefly, for de-lipidation, tissue slabs were placed in individual glass scintillation vials containing 20 mL of EZ Wash solution consisting of 50% (v/v) tetrahydrofuran prepared in sterile Milli-Q water. Vials were wrapped in foil for protection from light and shaken gently on a plate shaker inside a vented chemical fume hood for 16 hours at room temperature. To prepare the EZ View sample mounting and imaging solution, 100 g of 80% Nycodenz, 52.5 g of 7M urea, and 31.25 mg of 0.05% sodium azide powder were added to a 250 mL beaker. The beaker was placed on a hot plate, stirred using a magnetic stir bar, and gently heated to 37°C until the powders were fully dissolved. Aliquots (20 mL) of 0.02M sodium phosphate buffer (pH 7.4) were gradually added until the solution reached a total volume of 125 mL. The EZ View solution was left to mix overnight at room temperature. The next day, the EZ wash solution was removed from scintillation vials and sections were washed 4x with 20 mL Milli-Q water (1 hour per wash) to remove residual tetrahydrofuran. After the last wash, sections remained in a ventilated fume hood for an additional 10 minutes to evaporate residual water. Next, the EZ View solution was filtered through a vacuum filtration system to remove undissolved particulates. To render tissue sections transparent for two-photon imaging, tissue sections were then transferred to new glass scintillation vials and incubated in 5 mL of EZ View solution. Vials were wrapped in foil and gently shaken on a plate shaker at room temperature for 24 hours before two-photon imaging. Tissue slabs were imaged immediately after immunostaining in an EZ view-filled sample holder overlaid with a No. 0 coverslip. Two-photon imaging was performed with a 20-X, 1.0 NA objective lens (Olympus XLUMPLFLN 20XW), and sequential Z-stacks were collected for tdTomato/YFP (975 nm excitation) and anti-α-SMA antibody with Alexa fluor 647 conjugate (800 nm excitation).

### Hypoxyprobe injections and immunostaining

To measure brain-wide tissue hypoxia after chemogenetic stimulation of sparsely labeled pericytes, we performed Hypoxyprobe immunohistology. Hypoxyprobe (dissolved in sterile PBS) was retro-orbitally administered to tamoxifen-treated Atp13a5-Gq-DREADD mice at a dose of 60 mg/kg in a 50 μL volume under brief isoflurane anesthesia. Within 15-30 seconds of Hypoxyprobe injection, a 10 μg/kg bolus of DCZ or saline control was injected intraperitoneally. After awakening from anesthesia, mice were left in their home cages for 90 min for Hypoxyprobe binding and then trans-cardially perfused with PBS and 4% PFA. Brains were extracted and post-fixed overnight in 4% PFA in PBS with 0.02% sodium azide. The brains were then placed in 30% sucrose in PBS-azide for 2-3 days before embedding in OCT molds to cryoprotect brain specimens prior to sectioning.

Serially sliced histological sections (50 μm thickness) were collected with a cryostat, and then immunostained at five levels of the brain along the anteroposterior axis. Anterior sections were selected at two levels of the striatum, middle sections were selected at the levels of the dorsal and ventral hippocampal formation, and posterior sections of the cerebellum. Sections were washed 3x with PBS with 0.1% Triton-X and incubated in scintillation vials containing pab2627 rabbit antisera (1:100; Hypoxyprobe, #HP3-1000Kit) diluted in antibody solution (20% w/v Triton-X-100, 1% w/v normal goat serum, 0.01% w/v sodium azide and PBS). Scintillation vials were covered with aluminum foil and left overnight on a plate shaker. The next day, sections were washed 3x using PBS with 0.1% Triton-X before being placed in a secondary antibody solution containing goat-anti-rabbit Alexa Fluor 647 (1:500; Invitrogen, #A21244) in antibody solution for two hours at RT. Sections were then mounted in Fluoromount-G, cover slipped and left to dry in the dark overnight. All stained sections were imaged using a 12-bit camera on the Olympus APX100 slide scanner at 4x magnification. Optimal exposure times for Hypoxyprobe fluorescence detection (Ex: 670 nm; Em: 625 nm) was determined manually and kept consistent across all imaging sessions.

#### Brain region-of-interest (ROI) generation in ImageJ/Fiji

Slide scanned composite images were loaded into ImageJ/Fiji and fluorescent channels were split. Each section was matched to the Allen Brain Atlas coronal reference atlas and ROI contours were traced in ImageJ on a screen-mirrored tablet with a stylus. ROI contours were saved as .roi files for each section and loaded into a MATLAB script using the ReadImageJROI function, enabling ROIs for a tissue section to be imported into MATLAB without Java using ReadImageJROI.^52^ To ensure that ROI contours were identified in MATLAB, .roi files were saved in the same directory as the .TIFF sections and the ReadImageJROI function scripts to ensure proper calling of functions.

#### Hypoxyprobe quantification

With ROI contours generated, hypoxic microdomains within tissue were manually counted and collated across serial sections in a spreadsheet for each mouse. Hypoxic foci were defined as microdomains only if a contiguous region of immunolabeling exceeded 1,000 µm^2^. A Matlab script was then used to quantify hypoxia in each section. Briefly, pixel intensities bound within all ROI contours were stored in an array and the average pixel intensity value across five serial sections per animal was calculated. All pixel intensities bounded by ROI contours for a given section were normalized to the average pixel intensity value calculated for each mouse to allow better comparison across mouse cohorts. To summarize hypoxic microdomain burden across the brain, measurements were consolidated into nine higher-order anatomical ROI groups (**Fig. 3L**). At this level, the total number of microdomain was normalized to the sum of all ROI area measurements (in mm^2^) for ROIs belonging to an anatomical group. For example, the white matter anatomical group measurement reflects the total number of microdomains from the corpus callosum, fimbria, internal capsule, arbor vitae, and anterior commissure divided by the total sum of all ROI areas across all these regions. In total, the data reflected 519 distinct ROI measurements across DCZ treated mice (n = 6), and 378 measurements across saline controls (n = 4). To study the effects of treatment and anatomical ROI groups on hypoxic microdomain density, data was analyzed using two-way ANOVA with treatment (saline vs DCZ) and anatomical ROI groups as fixed factors. *Post-hoc* multiple-comparisons testing (Šídák’s correction) was used to compare microdomain density between DCZ- and saline-treated animals. In **Fig. 3L**, each datapoint reflects the hypoxic microdomain density for a certain anatomical ROI group for one mouse.

For analyses of **Fig. 3M**, somatosensory cortex microdomain area measurements were pooled across DCZ-treated mice (n = 6) and compared to convex-hull area estimations of blood-flow changes in pericyte contracted capillary segments *in vivo*. The convex hull area was bounded by distal branch-points of capillary segments experiencing ≥ 50% blood flow reduction from baseline. Somatosensory cortex microdomain areas were further binned based on size, and frequency counts for each size bin was plotted as the proportion of all bin measurements.

To establish the relationship between hypoxic microdomain density and capillary length density^2^ in **Fig. 3N**, respective data was fit to a second order decay quadratic model: y = B0 + B1x + B2x2. Fitted coefficients: B0 = 47.8 (95% CI [25.7 to 70.0]), B1 = -94.8 (95% CI [-149.8 to -39.8]), B2 = 47.8 (95% CI [14.4 to 81.3]). Parameters were estimated with least square regression without weighing, and 95% confidence intervals were computed using profile likelihood. Hypoxic microdomain density values were averaged across all Gq-DREADD-positive animals treated with DCZ (n = 6) and shown as individual data points. Capillary length density measurement for the insular piriform area reflects the mean of the agranular insular area and the piriform area; the septal complex measurement reflects the mean of the lateral and medial septal complex; the striatal-pallidal area measurement reflects the mean of the striatum and pallidum (all reported as mean without SEM).

### Magnetic Resonance Imaging

MRI was performed at the Oregon Health & Science University Advanced Imaging Research Center using a Bruker-Biospin 11.75T small animal MR system with a ParaVision 360 v3.6 software platform, and 10 cm inner diameter gradient set. Mice were anesthetized with a ketamine/xylazine mixture (5 mg xylazine/70 mg ketamine/kg), positioned with heads immobilized on an animal cradle and provided with 100% oxygen. Body temperature of the mice was monitored and maintained at 37°C while monitoring respiration (SA Instruments, Stony Brook, NY, United States). Mice were scanned employing a 72 mm (ID) 60 mm (length) Bruker RF resonator for transmitting and a Bruker 4 channel actively decoupled mouse head surface coil for receiving. For each mouse, a coronal 25-slice T_2_-weighted image was acquired (ParaVision spin echo RARE, 32 mm FOV to match the subsequent ASL images, 256×256 matrix, 125×125 µm in-plane resolution, 0.5 mm slice width, TR/TE 4000/38 ms, RARE factor 8, 2 averages). These T_2_-weighted anatomical scans were used for positioning the perfusion image slice at a consistent position centered over the hippocampus

Cerebral blood flow (ml/min/100 g) was measured using Arterial Spin Labeling (ASL) MRI, employing the flow-sensitive alternating inversion recovery rapid acquisition with relaxation enhancement pulse sequence (ParaVision FAIR-RARE, with TE/TR = 25/10000 ms, slice thickness 1 mm, number of slices = 1, matrix = 128×128, 250 µm in-plane resolution, RARE factor 32, 14 turbo inversion recovery values ranging from 20 to 4500 ms, and acquisition time of 14 min). This sequence labels the inflowing blood by global inversion of the equilibrium magnetization.^53^ Both the T2-weighted and ASL sequence were implemented at the baseline condition and a second time, starting 5 min after the injection of 10-35 µg/kg DCZ intraperitoneally to activate the Gq-DREADD.

#### ASL CBF data analysis

We implemented a custom preprocessing pipeline using FSL (v6.0)^46^ and ANTS (v2.4.4)^45^ to analyze CBF changes from ASL MRI data. T2-weighted images were bias field corrected using N4BiasFieldCorrection^54^ and brain extraction was performed using bet4animal, which included an initial center-of-mass estimation step to optimize the bet4animal parameters. Extracted brain masks were normalized (intensity-normalized to a mean of 1.0) to improve registration performance.

To determine regional CBF changes, brain masks were cropped, intensity-normalized and registered to a mouse anatomical atlas^55^, using a multi-stage (rigid, affine, SyN) nonlinear registration using antsRegistration. Atlas label maps were then transformed into subject space using antsApplyTransforms^56^ with nearest-neighbor interpolation.

For ASL quantification, the atlas label maps from the two contiguous anatomical slices overlapping the ASL slice, were brought into ASL space. For each of the two slices, region-specific masks were created for structures of interest including the hippocampus, cortex, amygdala, thalamus, hypothalamus, and ventricles. Masks were combined across the two slices using intersection logic for the non-ventricle regions (e.g., hippocampus, amygdala) or union logic for the ventricles. This way, perfusion in the thick ASL slice would only be considered to belong to a particular region if the voxel was labeled as the same region in both overlapping anatomical slices. When calculating global brain perfusion, voxels that were part of the ventricular system were removed in either of the overlapping anatomical images to reduce partial volume effects.

Perfusion values were extracted from the final ASL images using voxelwise intensity measures within each anatomical mask. Whole-brain perfusion estimates were derived after additionally thresholding voxel intensities to exclude extreme outliers (>3 standard deviations from the mean). Mean perfusion values for each region-of-interest were compiled for statistical analysis.

#### Statistics for MRI data

All statistical analyses were performed in R (version 4.4.1) using the lme4 and lmerTest packages. To account for the multiple regions for each mouse, mixed-effects linear models were used with mouse identity included as a random intercept term. This approach appropriately models the correlation among regional measures obtained from the same animal.

For analyses of cerebrovascular reactivity, the CBF following DCZ administration was modeled as a function of genotype (Group), brain region (Region), and their interaction, while controlling for baseline CBF as a covariate:

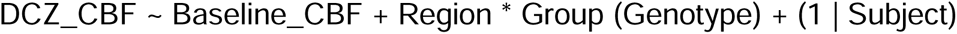

For analyses of baseline CBF, the following model was used:

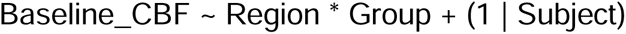

Significance of fixed effects was assessed using Type III ANOVA with Satterthwaite’s approximation for denominator degrees of freedom. Where main effects of region were significant, Tukey-adjusted post hoc comparisons were performed using estimated marginal means (emmeans package).

## Notes

### Competing Interest Statement

The authors have declared no competing interest.

