## Supplementary Figures for "Chemogenetic Pericyte Activation Reveals Broad Contractile Ability and Limbic Vulnerability to Capillary Flow Deficits"

[1] Center for Developmental Biology and Regenerative Medicine, Seattle Children's Research Institute, Seattle, WA, USA
[2] Norcliffe Foundation Center for Integrative Brain Research, Seattle Children's Research Institute, Seattle, WA, USA
[3] Department of Pediatrics, University of Washington, Seattle, WA
[4] Department of Bioengineering, University of Washington, Seattle, WA
[5] Department of Neurology, University of Washington, Seattle WA, USA
[6] Graduate Program in Neuroscience, University of Washington, Seattle, WA, USA
[7] Department of Anesthesiology, University of Colorado, Anschutz Medical Campus, CO, USA
[8] Department of Neurology Jungers Center for Neurosciences Research, Oregon Health & Science University, Portland, OR
[9] Helfgott Research Institute, National University of Natural Medicine, Portland, OR, USA
[10] Department of Neurosciences, Cleveland Clinic Research, Cleveland, OH 44106, USA
[11] Department of Molecular Medicine, Cleveland Clinic Lerner College of Medicine, Case Western Reserve University, Cleveland, OH, USA
[12] Zilkha Neurogenetic Institute, University of Southern California, Los Angeles, CA
[#] These authors contributed equally

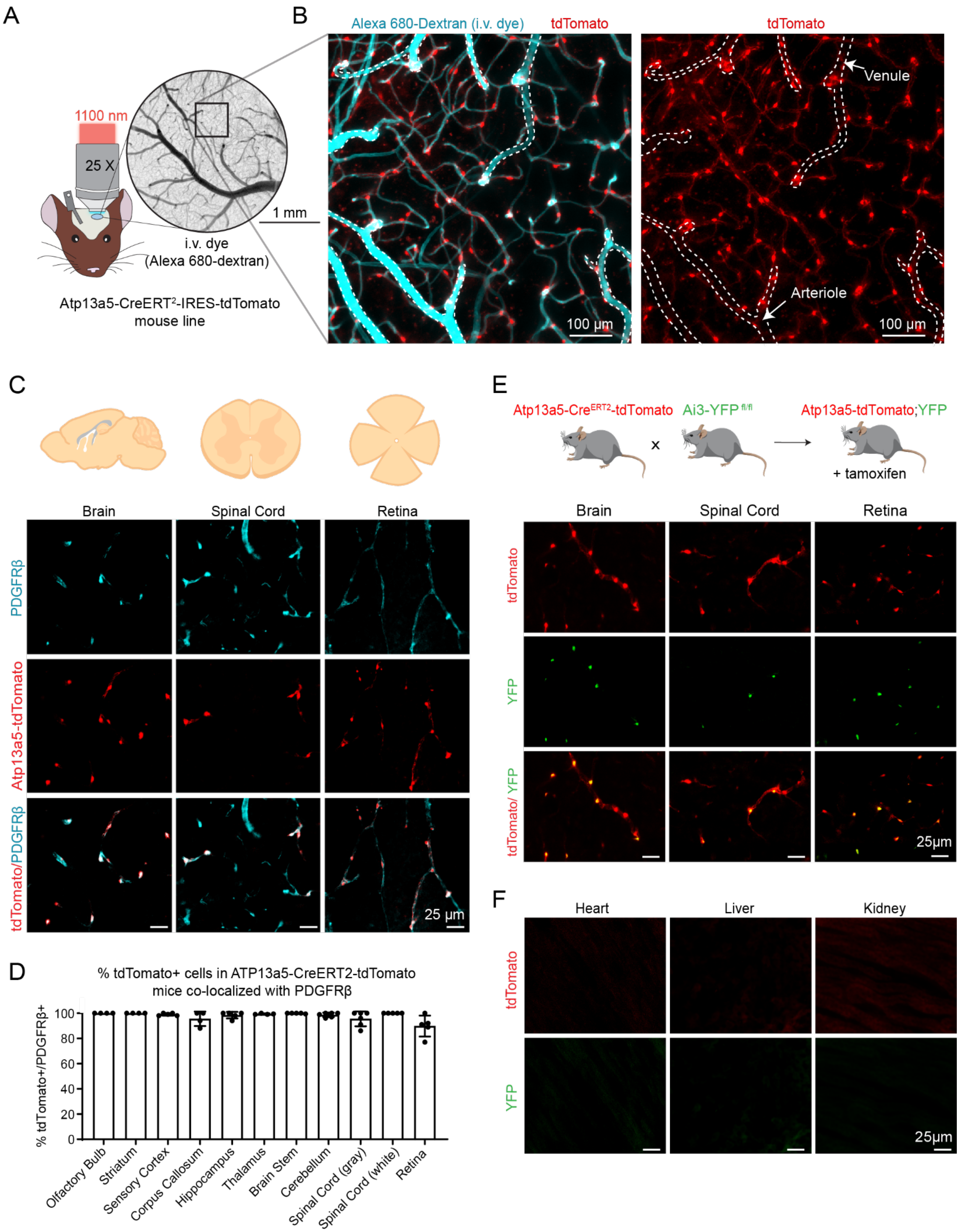
A
1100 nm
25 X
1 mm
i.v. dye
(Alexa 680-dextran)
Atp13a5-CreERT2-IRES-tdTomato
mouse line
B
Alexa 680-Dextran (i.v. dye)
tdTomato
Venule
Arteriole
100 µm
C
Brain
Spinal Cord
Retina
PDGFRβ
Atp13a5-tdTomato
tdTomato/PDGFRβ
25 µm
D
% tdTomato+ cells in ATP13a5-CreERT2-tdTomato mice co-localized with PDGFRβ
% tdTomato+/PDGFRβ+
100
80
60
40
20
0
Olfactory Bulb
Striatum
Sensory Cortex
Corpus Callosum
Hippocampus
Thalamus
Brain Stem
Cerebellum
Spinal Cord (gray)
Spinal Cord (white)
Retina
E
Atp13a5-CreERT2-tdTomato
Ai3-YFP fl/fl
Atp13a5-tdTomato;YFP
x
+ tamoxifen
Brain
Spinal Cord
Retina
tdTomato
YFP
tdTomato/ YFP
25µm
F
Heart
Liver
Kidney
tdTomato
YFP
25µm

**Supplementary Figure 1. Characterization of tdTomato expression in Atp13a5-CreERT2 mice in the central nervous system and peripheral organs.**

**(A)** Schematic diagram of *in vivo* two-photon imaging set up with a cranial window in a Atp13a5-CreERT2 mouse, with Alexa 680-Dextran labeled blood vessels used as landmarks. The inset in the cranial window shows the region imaged in B.

**(B)** Representative 100 μm thick average projection of Atp13a5-tdTomato-positive pericytes (red) and Alexa 680-Dextran-labeled vessel lumen (cyan) in layers II/III of the sensory cortex of a 9-month-old Atp13a5-CreERT2 mouse using *in vivo* multi-photon imaging.

**(C)** Representative images of PDGFRβ (cyan) and tdTomato (red) positive cells and co-localization in the brain, spinal cord and retina of Atp13a5-CreERT2.

**(D)** Percentage of tdTomato-positive cells colocalized with PDGFRβ-positive cells in different CNS regions in 4–5-month-old Atp13a5-CreERT2;Ai3 mice (n=4-6 female mice).

**(E)** Schematic of cross-breeding to generate Atp13a5-CreERT2;Ai3 mice. Representative images of tdTomato (red) positive cells expressing endogenous YFP (green) in the brain, spinal cord and retina of a 3-month-old Atp13a5-CreERT2;Ai3 mouse, 1-month post-tamoxifen treatment.

**(F)** Representative images highlighting the absence of tdTomato (red) and YFP (green) positive cells in the heart, liver and kidney in a 4-month-old Atp13a5-CreERT2;Ai3 mouse.

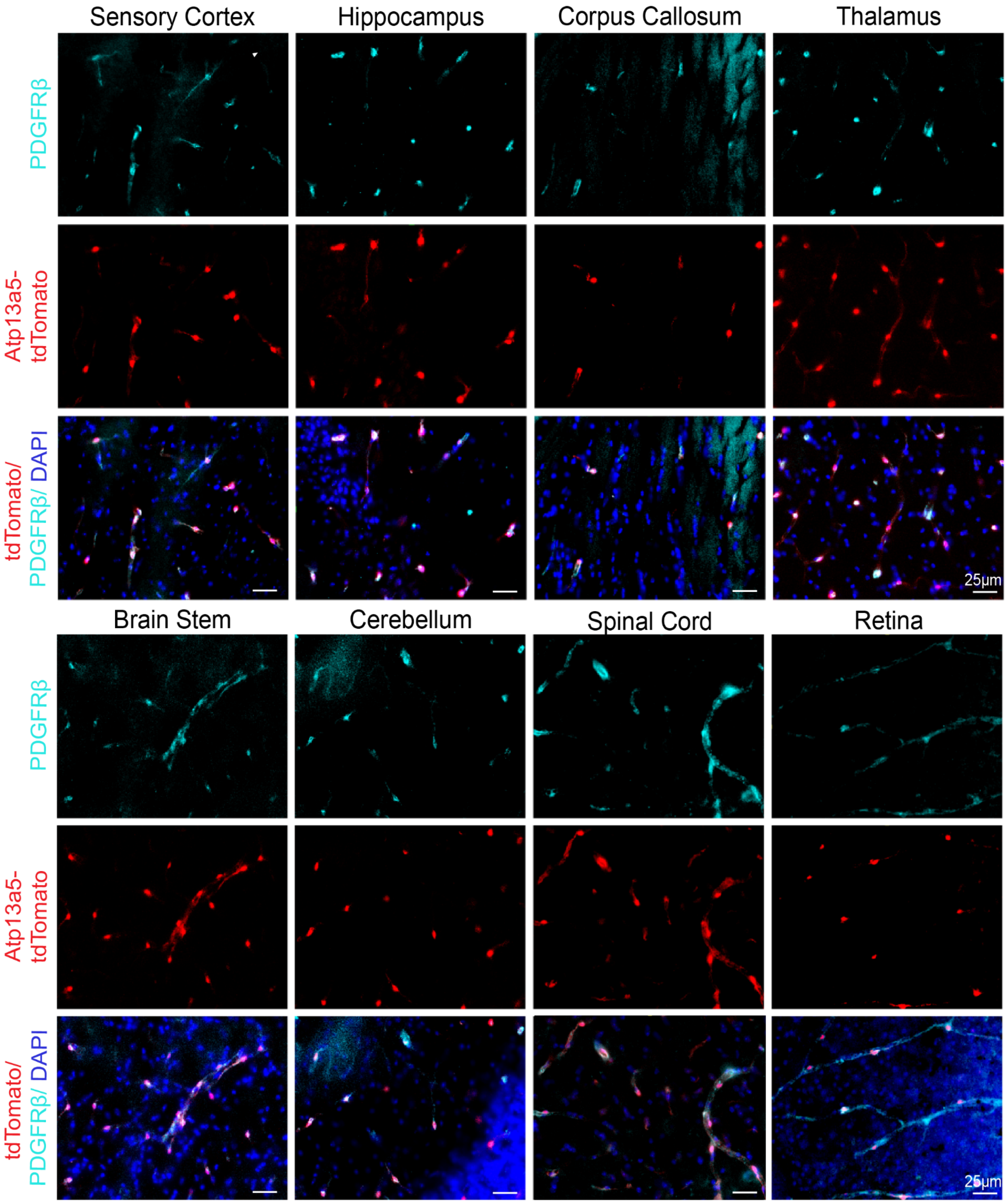


**Supplementary Figure 2. Endogenous tdTomato expression across different regions of the central nervous system in Atp13a5-CreERT2 mice.** Representative images of PDGFRβ (cyan), tdTomato (red) and DAPI (blue) positive cells across different regions of the brain, spinal cord and retina in a 4-month-old Atp13a5-CreERT2;Ai3 mouse. Images highlight co-localization of PDGFRβ and Atp13a5-tdTomato.

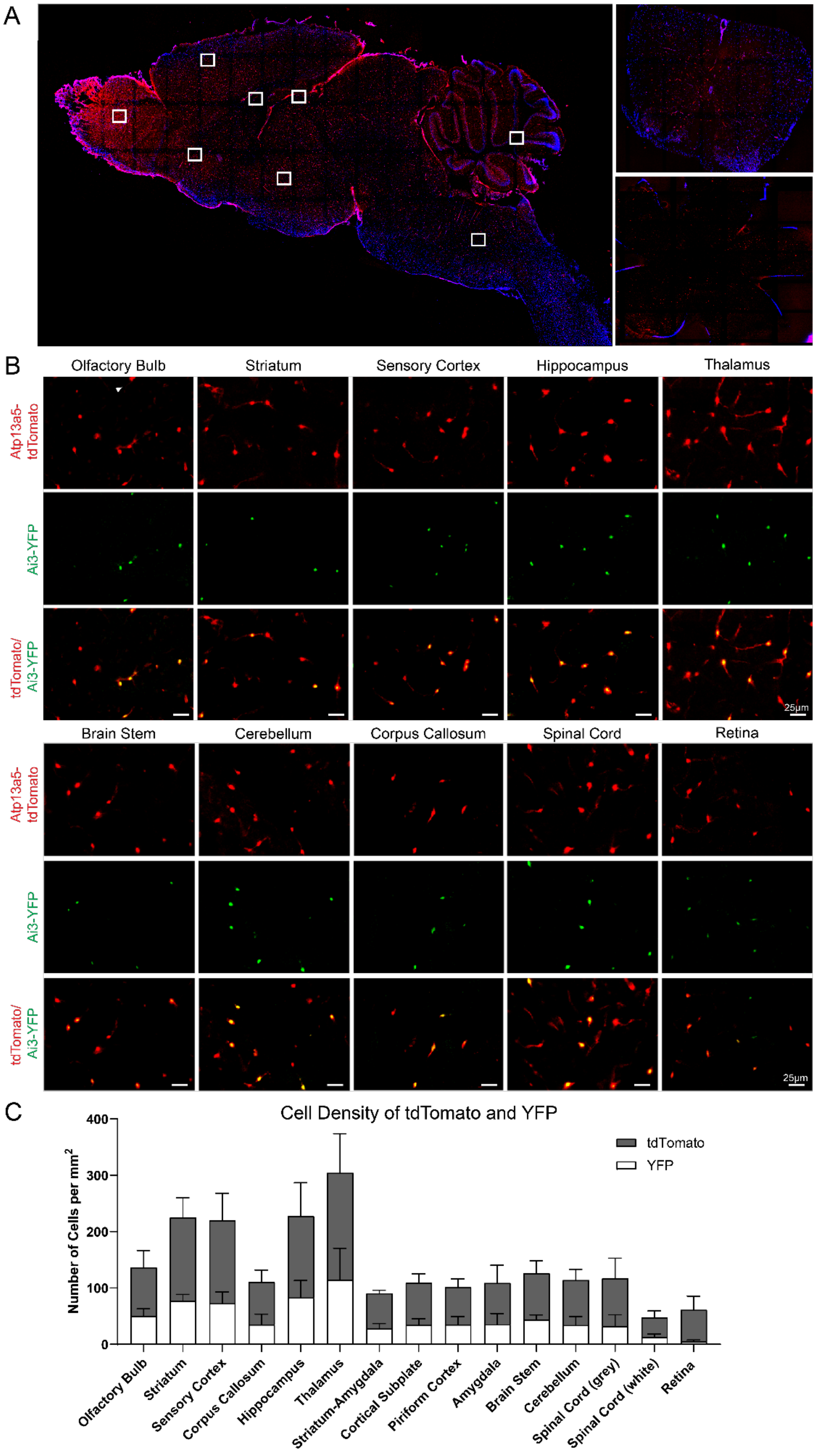
A
B
Olfactory Bulb
Striatum
Sensory Cortex
Hippocampus
Thalamus
Atp13a5-tdTomato
Ai3-YFP
tdTomato/Ai3-YFP
25μm
Brain Stem
Cerebellum
Corpus Callosum
Spinal Cord
Retina
C
Cell Density of tdTomato and YFP
Number of Cells per mm²
400
300
200
100
0
tdTomato
YFP
Olfactory Bulb
Striatum
Sensory Cortex
Corpus Callosum
Hippocampus
Thalamus
Striatum-Amygdala
Cortical Subplate
Piriform Cortex
Amygdala
Brain Stem
Cerebellum
Spinal Cord (grey)
Spinal Cord (white)
Retina

**Supplementary Figure 3. Endogenous tdTomato and tamoxifen-induced YFP expression in different regions of the central nervous system in Atp13a5-CreERT2;Ai3 mice.**

**(A)** Representative image of endogenous tdTomato expression (red) and DAPI (blue) in a sagittal cut brain, coronal spinal cord (lumbar) and whole mount retina of a 5-month Atp13a5-CreERT2;Ai3 mouse. Insets on brain image are regions of the brain in B.

**(B)** Representative images of endogenous tdTomato-positive cells (red) expressing tamoxifen-induced YFP (green) across different regions of the brain, spinal cord and retina in a 4-month-old Atp13a5-CreERT2;Ai3 mouse, 1-month post-tamoxifen treatment.

**(C)** Stacked bar graph showing number of tdTomato- (in gray) and YFP-positive cells (in white) per $mm^2$ in different regions of the brain, spinal cord and retina in 4–5-month-old Atp13a5-CreERT2;Ai3 mice (n=3-6 female mice).

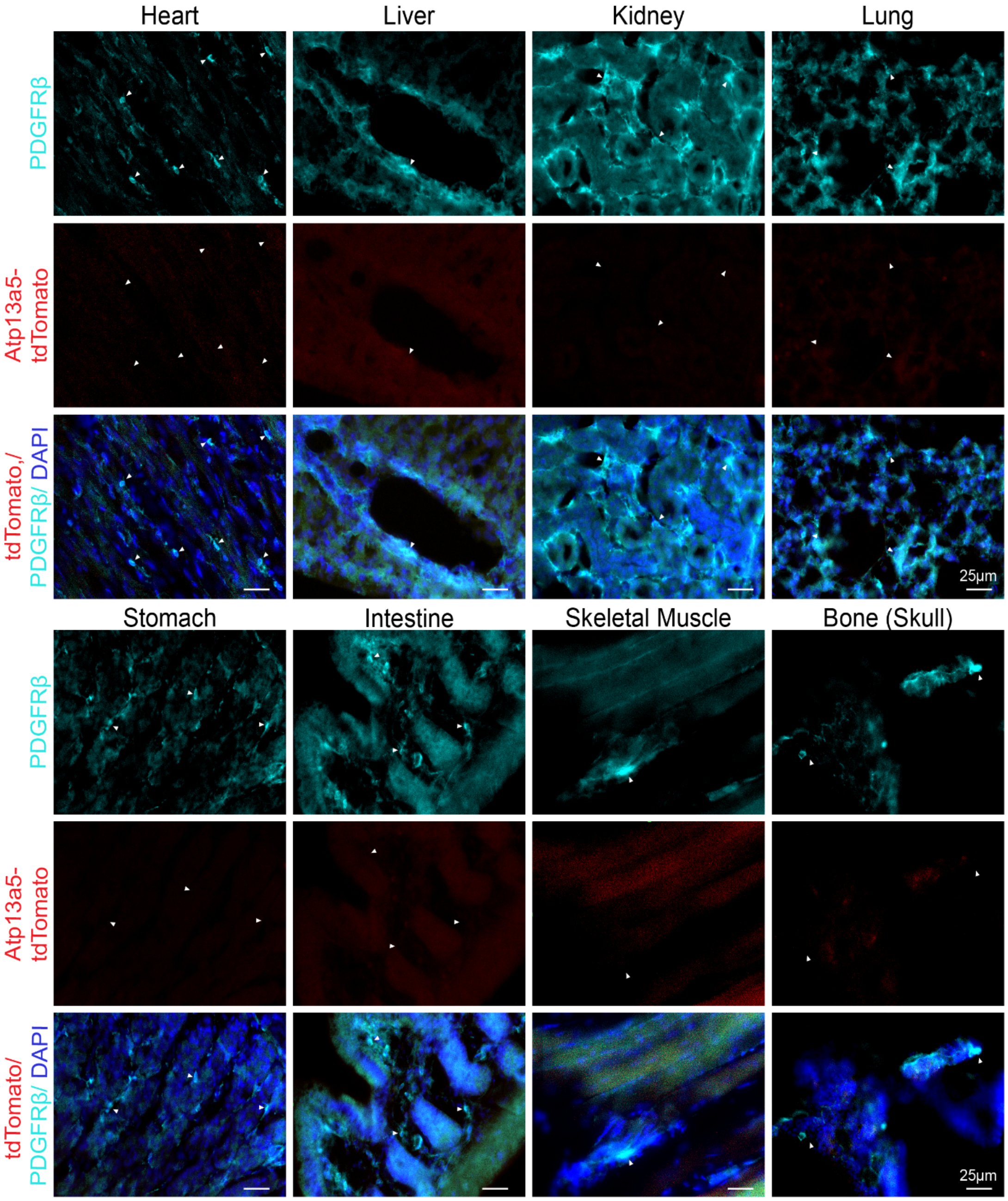


**Supplementary Figure 4. Endogenous tdTomato expression is absent in peripheral organs in Atp13a5-CreERT2 mice.** Representative images of PDGFRβ (cyan), tdTomato (red) and DAPI (blue) positive cells in the heart (ventricle), liver, kidney, lung, stomach, intestine (duodenum), skeletal muscle (gastrocnemius) and bone (skull) in a 5-month-old Atp13a5-CreERT2 mouse. White arrows highlight the absence of co-localization of tdTomato with PDGFRβ-positive cells.

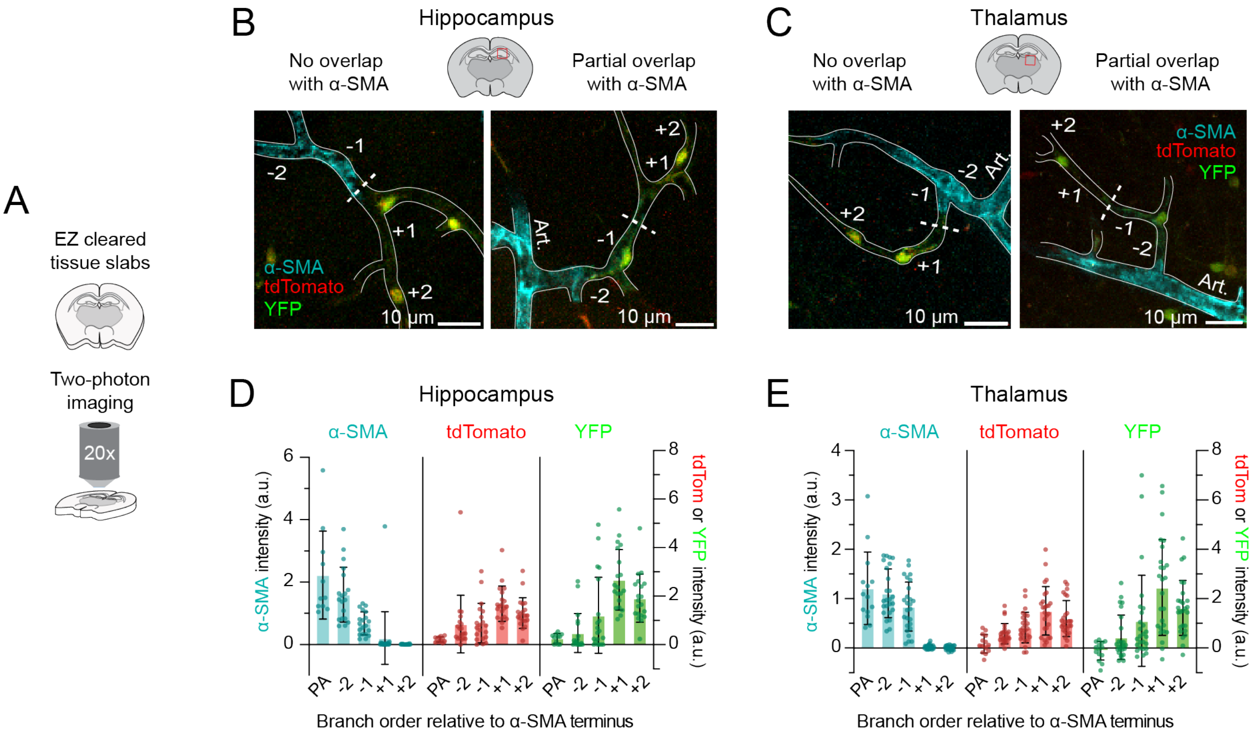


**Supplementary Figure 5. Genetic targeting in the ACT zone of hippocampus and thalamus in Atp13a5-CreERT2;Ai3 mice.**

**(A)** Optically-cleared brain tissue slabs immunostained for α-SMA and imaged by two-photon to evaluate fluorescent protein expression in relation to the ACT zone.

**(B,C)** Fluorescent reporter expression as a function of branch order from α-SMA terminus (dotted line).

**(D,E)** Bar graph showing reporter intensity at the penetrating arteriole (PA) and different branch orders from the α-SMA terminus. Negative branch orders correspond to the ACT zone, covered by α-SMA-positive ensheathing pericytes. Positive branch orders correspond to the capillary zone, covered by α-SMA-low/undetectable capillary pericytes.

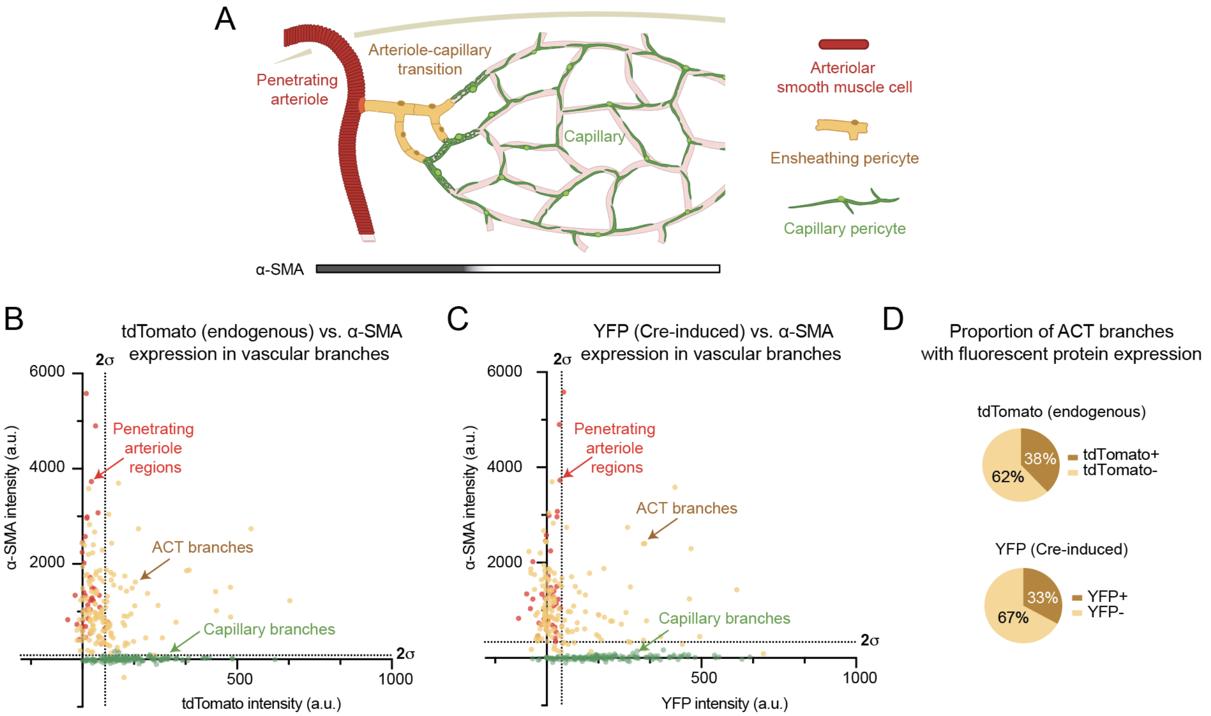


**Supplementary Figure 6. Quantification of genetic targeting in different vascular zones of cortex in Atp13a5-CreERT2;Ai3 mice.**

**(A)** Schematic showing penetrating arteriole, ACT, and capillary zones. Penetrating arteriole and ACT zones express α-SMA, but the capillary zone expresses low to undetectable levels.

**(B)** Scatterplot showing α-SMA immunolabeling intensity plotted as a function of endogenous tdtomato intensity for each microvessel branch sampled, including penetrating arteriole regions (red), ACT branches (yellow), and capillary branches (green). Dotted lines show the mean + 2σ threshold for penetrating arterioles (vertical dotted line) and capillary branches (horizontal dotted line).

**(C)** Scatterplot showing α-SMA immunolabeling intensity plotted as a function of tamoxifen-induced YFP expression intensity. Dotted lines show the mean + 2σ threshold for penetrating arterioles (vertical dotted line) and capillary branches (horizontal dotted line).

**(D)** Proportion of total ACT zone branches with fluorescence intensities outside the + 2σ thresholds in panels B,C. That is, branches that are significant for both α-SMA and endogenous tdTomato (upper panel), or both α-SMA and tamoxifen-induced YFP (lower panel).

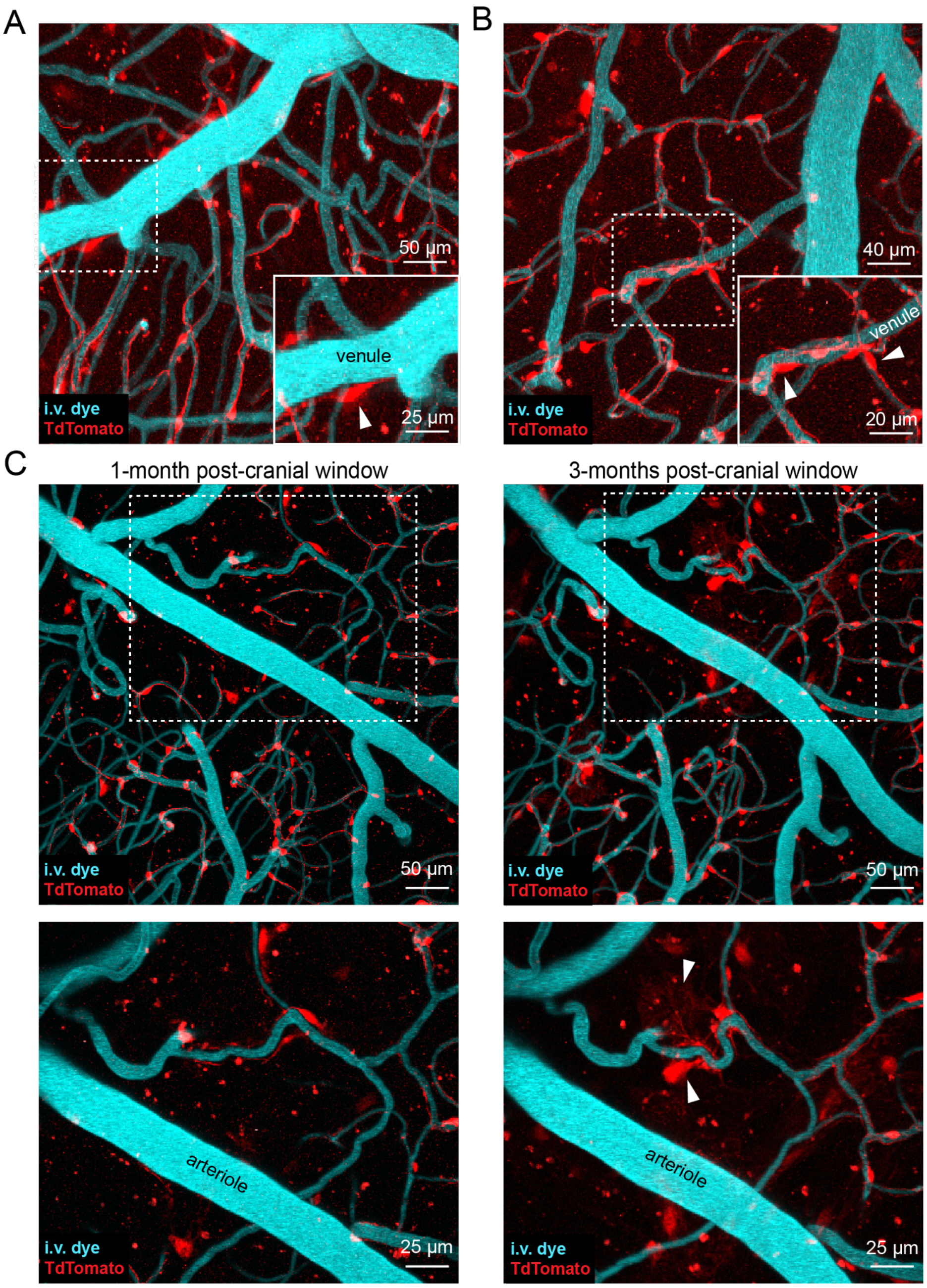


**Supplementary Figure 7. TdTomato expression in perivascular cells of pial and ascending venules, and sparse labeling of fibroblast-like cells in the meninges after cranial window implantation in Atp13a5-CreERT2 mice.**

**(A,B)** Representative 50 µm thick average projections of tdTomato-positive cells (red) and Alexa 680-Dextran-labeled vessels (i.v. dye; cyan) in layer I of the sensory cortex of a 9-month-old Atp13a5-CreERT2 mouse collected using *in vivo* multi-photon imaging. TdTomato-positive mural cells (arrowheads) found on pial venule (A) and superficial region of ascending venule (B).

**(C)** Potential tdTomato-positive fibroblasts (arrowheads) in meninges absent at 1-month post-cranial window implantation but present at 3-months.

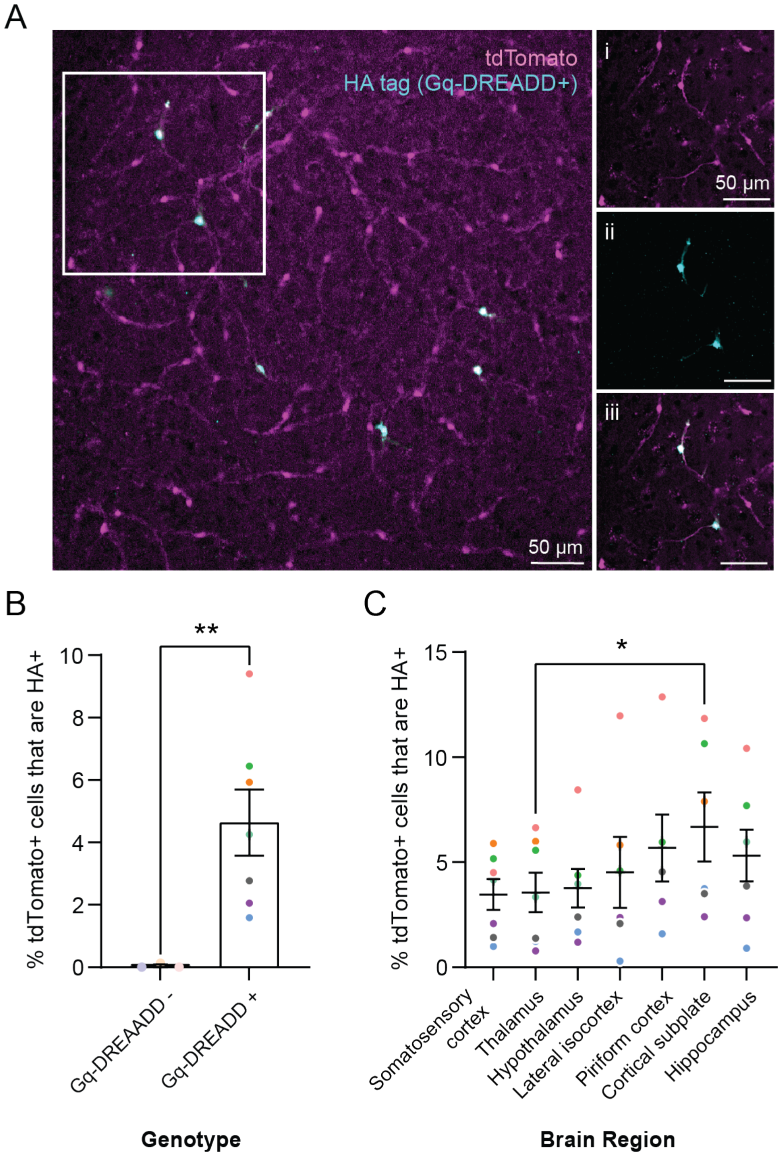


**Supplementary Figure 8. Genetic recombination rate in Atp13a5-Gq-DREADD.**
(**A**) Gq-DREADD-expressing pericytes in tamoxifen-injected Atp13a5-Gq-DREADD are identified by colocalization of endogenous tdTomato (magenta) and HA tag (cyan) positivity. Insets show tdTomato (i), HA-tag (ii) and overlay of these channels (iii).
(**B**) Recombination rate in Atp13a5-Gq-DREADD mice is variable across mice (symbol color) and occurs in 4.64 ± 1.06% (mean ± s.e.m; n = 7) of tdTomato-positive pericytes, in contrast to Gq-DREADD-negative littermates that do not show HA-tag positivity above background (p = 0.0049, unpaired t-test with Welch's correction).
(**C**) Gq-DREADD recombination rate varies by brain region in Gq-DREADD-positive mice (p = 0.0419, n = 7, mixed effects analysis with Geisser-Greenhouse correction). *Post-hoc* tests indicated the cortical subplate had modestly higher recombination rate than the thalamus (p = 0.0435, n = 7, Tukey's multiple comparisons test).

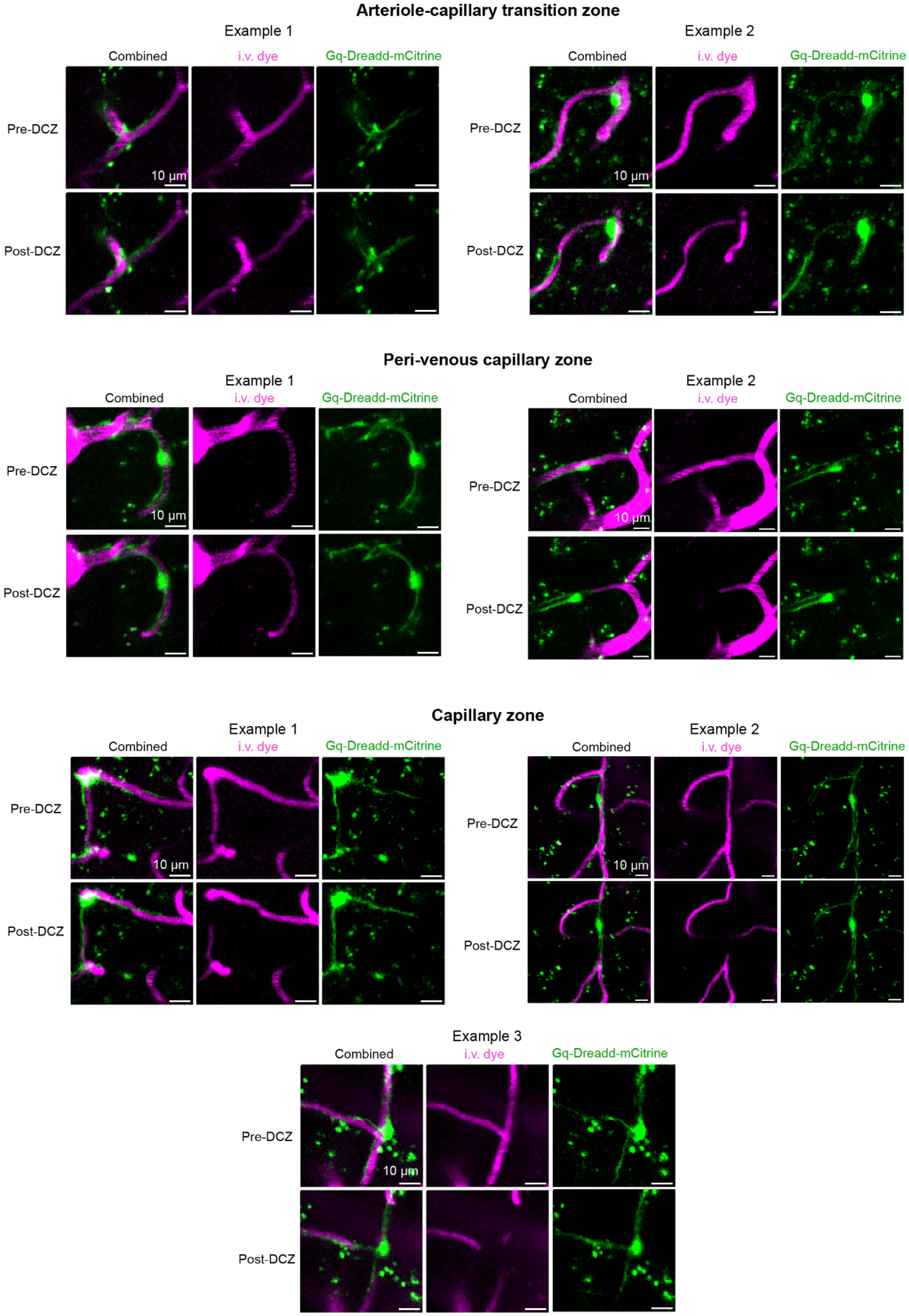


**Supplementary Figure 9. Additional examples of chemogenetically-induced pericyte contraction.** Examples are shown from ACT, peri-venous capillary, and capillary zones with overlays of mCitrine-fused Gq-DREADD and i.v. dye, and each channel shown separately.

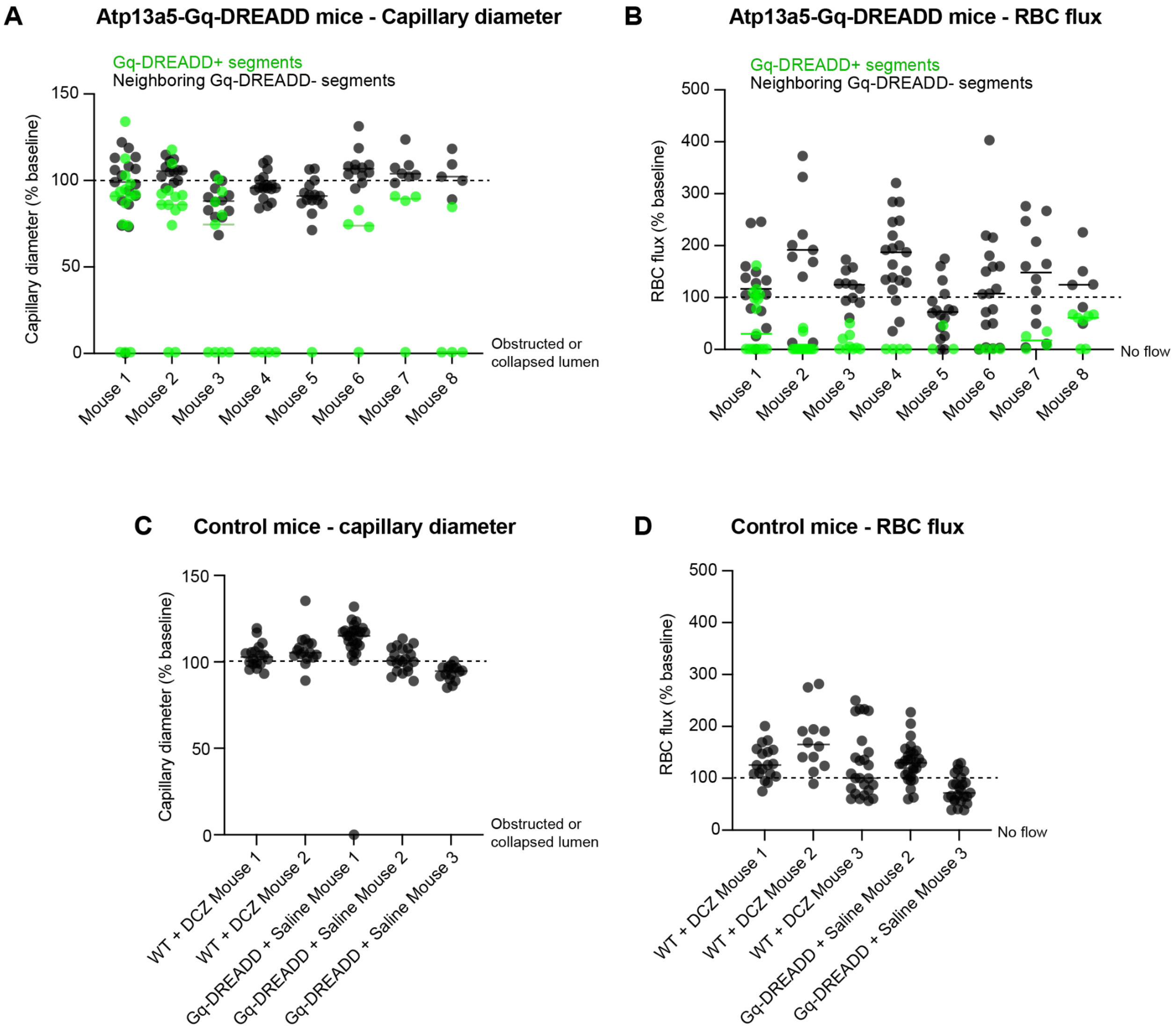


**Supplementary Figure 10. Consistency of chemogenetic pericyte contraction across Atp13a5-Gq-DREADD mice and absence of contraction in controls.**

**(A,B)** Capillary diameter and RBC flux change from each Atp13a5-Gq-DREADD mouse within Figure 2G,H. **(C,D)** Capillary diameter and RBC flux change from each control mouse within Figure 2G,H. Controls consistent of WT littermates (Atp13a5-CreERT2-tdTomato-positive but Gq-DREADD negative) treated with 10 mg/kg DCZ, or Atp13a5-Gq-DREADD mice treated with saline. Capillary diameter for one WT+DCZ mouse, and RBC flux for one Atp13a5-Gq-DREADD+saline mouse was not analyzed due to insufficient quality of images and/or line-scans.

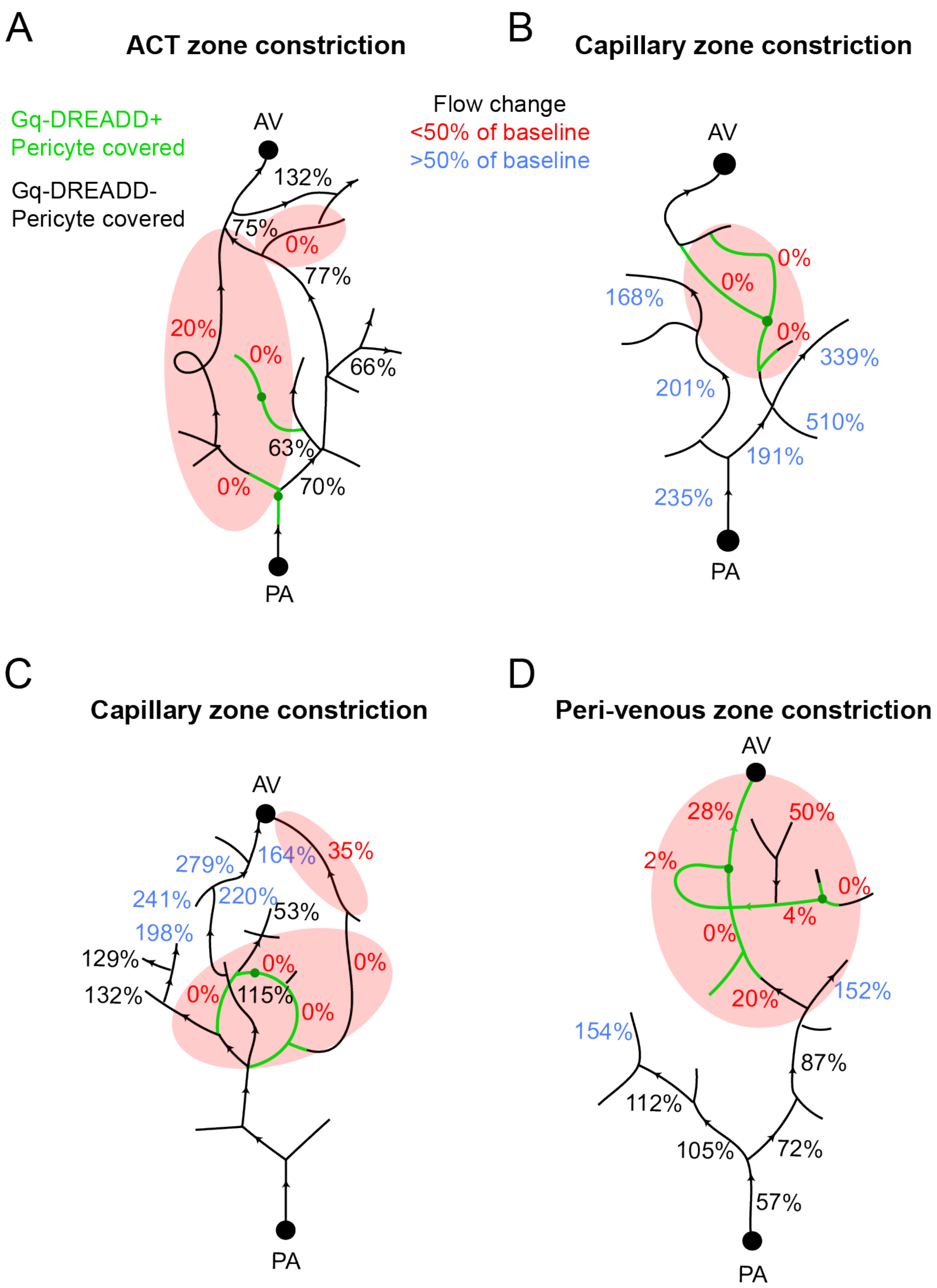


**Supplementary Figure 11. Qualitative assessment of blood flow across the capillary network after pericyte contraction in different microvascular zones.**

**(A)** Example of ensheathing pericyte contraction, and nearby capillary pericyte contraction, in the ACT zone of a penetrating arteriole (PA) offshoot, causing a broader region of hypoperfusion.

**(B, C)** Two examples of isolated pericyte contraction in the mid-capillary zone, associated with a local hypoperfusion, but also hyperperfusion in neighboring regions.

**(D)** Example of pericyte contractions (two adjacent Gq-DREADD-positive cells) near an ascending venule (AV) leading to a broader region of hypoperfusion.

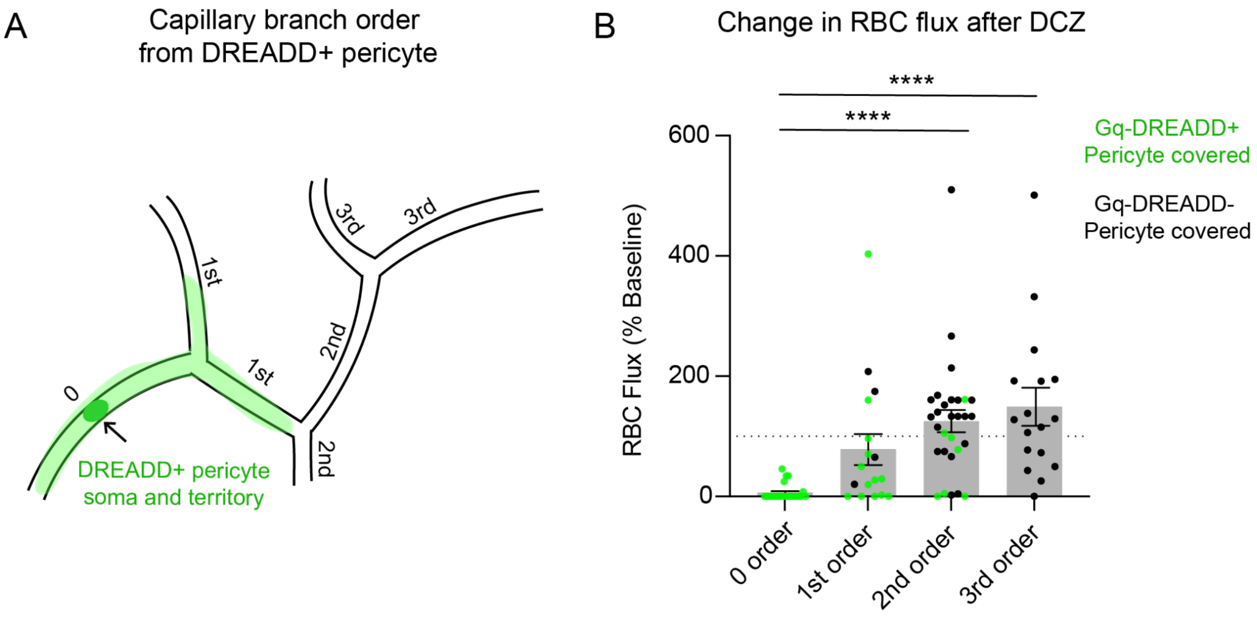


**Supplementary Figure 12. Reduction of RBC flux is greatest near the pericyte soma. (A)** Schematic diagram showing method for capillary branch ordering to be used in panel B. **(B)** Change in RBC flux from baseline after DCZ injection to Atp13a5-Gq-DREADD mice. Graph shows effect at different capillary branch orders from capillary segment in contact with the Gq-DREADD+ pericyte soma (denoted 0 order). ****p<0.001, two-way ANOVA with bonferroni *post-hoc* test.

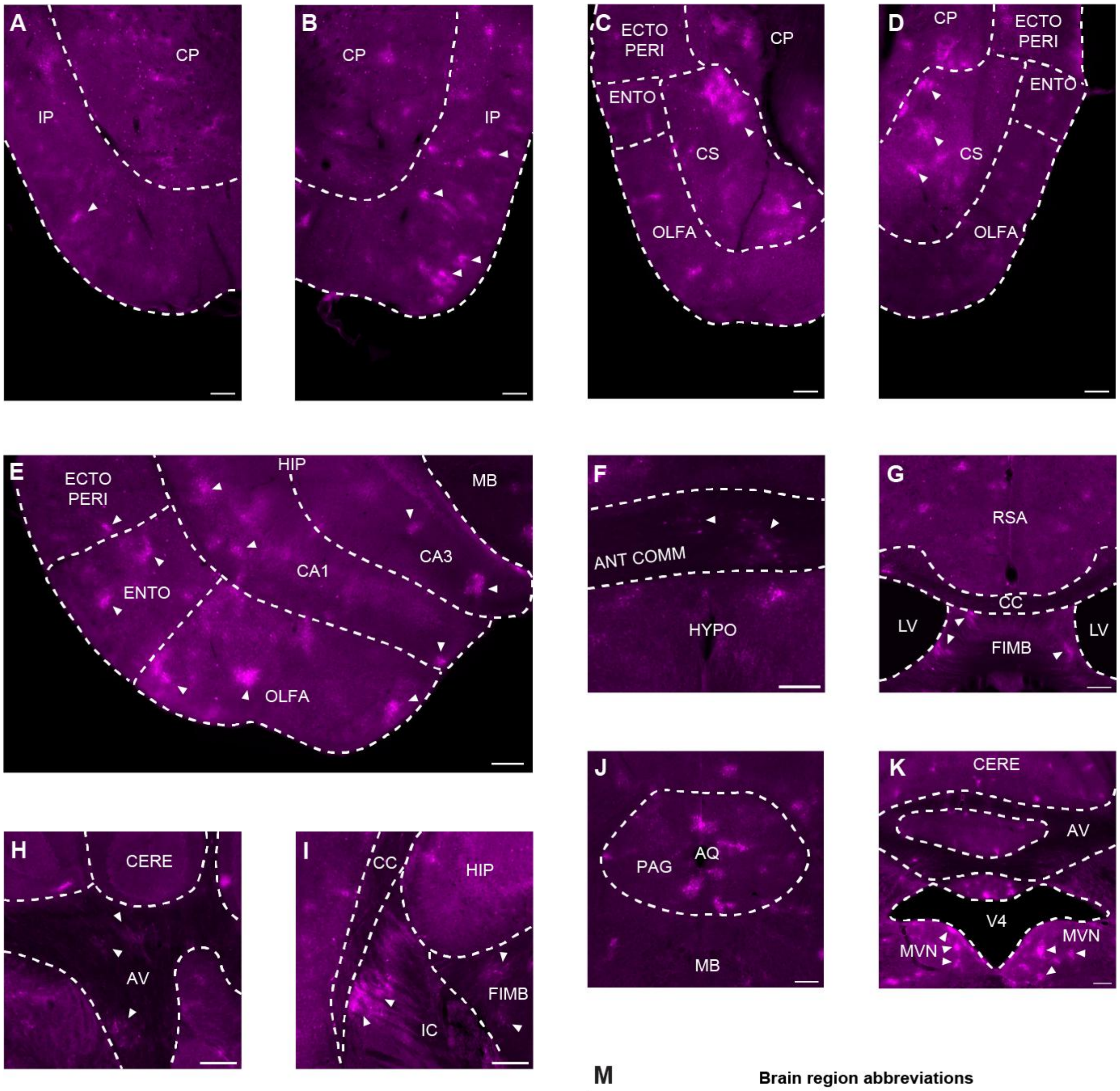


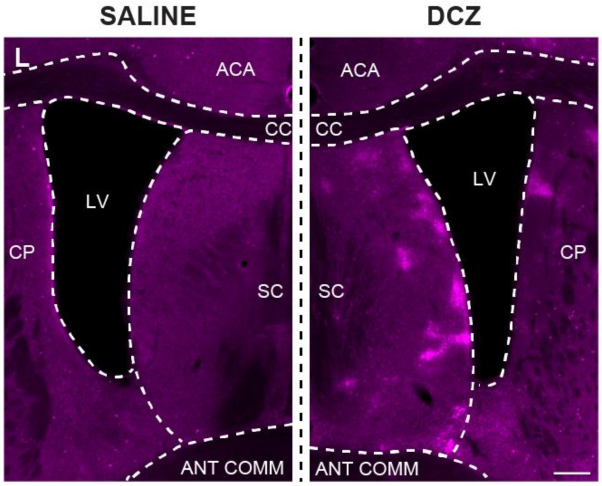


**M** **Brain region abbreviations**

ACC: anterior cingulate cortex
ANT COMM: anterior commissure
AQ: cerebral aqueduct
AV: arbor vitae (cerebellum)
CA1: Ammon's horn - field CA1
CA3: Ammon's horn - field CA3
CC: corpus callosum
CERE: cerebellum
CP: caudate-putamen
CS: cortical subplate
ECTO: ectorhinal assoc. cortex
ENTO: entorhinal cortex
FIMB: fimbria
HIP: hippocampal formation
HYPO: hypothalamus
IC: internal capsule
IP: insular-piriform area
LV: lateral ventricle
MB: midbrain
MVN: medial vestibular nucleus
OLFA: olfactory area
PERI: perirhinal assoc. cortex
RSA: retrosplenial area
SC: septal complex
V4: 4th ventricle

**Supplementary Figure 13. Hypoxic microdomains in Atp13a5-Gq-DREADD mice.**

**(A,B)** Hypoxyprobe staining within sections containing the insular and piriform cortices.

**(C-E)** Hypoxic micropockets in the cortical subplate (C,D), and the hippocampus and olfactory area (E).

**(F-I)** Hypoxyprobe staining in the white matter, including anterior commissure (F), corpus callosum and fimbria (G), arbor vitae of the cerebellum (H), and between fiber tracts of the corpus callosum and internal capsule (I).

**(J,K)** Hypoxic microdomains in the periaqueductal grey of the midbrain (J) and the medial vestibular nucleus of the cerebellum (K).

**(L)** Comparison of labeling in the septal complex in saline-controls (left) and DCZ-treated (right) Atp13a5-Gq-DREADD brains. Scale for panels A-L: 200 µm.

**(M) L**egend of brain region abbreviations.

### Supplementary Movie Legends

**Supplementary Movie 1.** Example of Gq-DREADD+ thin-strand pericyte contracting within peri-venous zone. Note heightened contraction near the pericyte soma, and formation of a divot when the thin process wraps around a capillary above the soma. The pericyte processes also pull the capillary longitudinally and distort the endothelium. Shown in Fig. 2I.

**Supplementary Movie 2.** Example of Gq-DREADD+ thin-strand pericyte contracting within peri-venous zone. Note heightened contraction near the pericyte soma, but constriction of the capillary is also apparent near a branch of the process.

**Supplementary Movie 3.** Example of Gq-DREADD+ thin-strand pericyte contracting near the venous zone. Thin pericyte processes (soma out of view) contact capillaries 1-2 branch orders away from a venule and exhibit robust contraction. The processes pull the capillary longitudinally. Shown in Fig. 2F.

**Supplementary Movie 4.** Example of Gq-DREADD+ thin-strand pericyte contracting within the mid-capillary zone. Thin processes above the soma constrict the capillary lumen. A process below the soma pulls longitudinally and distorts the underlying endothelium. Shown in Fig. 2K.

**Supplementary Movie 5.** Example of Gq-DREADD+ thin-strand pericyte contraction within mid-capillary zone. The thin processes constrict the capillary lumen and pull the capillary longitudinally to distort the endothelium.

**Supplementary Movie 6.** Example of Gq-DREADD+ thin-strand pericyte contraction within mid-capillary zone. A capillary segment above the pericyte soma transiently disappears, likely due to obstruction by a lodged blood cell during constriction of the lumen. Longitudinal pulling of the process below the soma distorts the endothelium.

**Supplementary Movie 7.** Example of Gq-DREADD+ thin-strand pericyte and putative mesh pericyte (defined only by morphology) contraction co-occurring in the same local network. Constrictions are seen specifically at locations contacted by the Gq-DREADD+ pericytes, which is particularly apparent with the mesh pericyte. Longitudinal pulling of the thin-strand process distorts the endothelium.
